# Centriolar satellites assemble via a hierarchical pathway driven by PCM1 multimerization

**DOI:** 10.1101/2025.07.27.666075

**Authors:** Efe Begar, Ece Seyrek, Selin Yilmaz-Karaoglu, Melis D. Arslanhan, Ezgi Odabasi, Elif Nur Firat-Karalar

## Abstract

Centriolar satellites (CS) are ubiquitous, membrane-less organelles recognized for organelle crosstalk, plasticity, diverse functions and links to developmental and neuronal diseases. However, the molecular principles governing their assembly and regulation remain poorly understood. To address this, we developed cellular and *in vitro* biogenesis assays that allow spatiotemporal quantification of CS granule properties during assembly, remodeling and maintenance. Using these tools, we show that CS assemble via a hierarchical pathway initiated by PCM1 scaffold formation followed by regulated client recruitment. PCM1 intrinsically assembles into granules through multimerization, a process modulated by cytoskeleton. High-resolution imaging revealed that PCM1 and its clients occupy distinct subdomains with different compositions and dynamics, adding an additional layer of regulation. Perturbing PCM1 multimerization impaired ciliary signaling, underscoring its functional importance. Collectively, these findings define the molecular basis of CS biogenesis, establish new tools to probe their context-dependent functions, and provide a framework for understanding how CS deregulation contributes to disease. More broadly, the principles uncovered here may extend to other membrane-less organelles, explaining their specificity and plasticity.

## Introduction

The dynamic organization of the cytoplasm by partitioning molecules into membrane-less organelles is an emerging mechanism for spatiotemporal control of cellular functions essential for development and physiology ^1–4^. Organelles such as stress granules, P-bodies, and germ granules form when scaffolding proteins multimerize, often coupled with phase separation, to create structures that concentrate macromolecules required for their functions ^5, 6^. These organelles rapidly assemble, disassemble and remodel in response to intrinsic and extrinsic stimuli, enabling cells to sense and adapt to changes in their environment. However, absence of a delimiting membrane poses challenges in regulating size, composition, and dynamics, and consequently, their functions^6^.

Centriolar satellites (CS) are ubiquitous, membrane-less organelles marked by their scaffolding protein Pericentriolar Material-1 (PCM1), which is essential for their assembly ^7–11^. They were first discovered in epithelial cells as ∼100 nm granules that cluster and move around the centrosomes in a microtubule-dependent manner ^9^. In these cells, PCM1 interacts with over 200 proteins, collectively referred to as clients, some of which are mutated in developmental and neurodegenerative disorders ^12–15^. CS function as part of a membrane-less trafficking pathway that stores, modifies and transports proteins to cellular structures such as centrosomes, cilia and autophagosomes ^8, 10–12, 16, 17^. Through these activities, CS regulate diverse cellular processes, many of which depend on this organelle crosstalk, and require highly dynamic and adaptive behavior ^7, 18–20^. Among these processes are cell signaling (e.g., Hedgehog), cell division, motility, stress response, proteostasis and neurogenesis, as revealed by loss-of function studies in cells and animal models ^7, 8, 10–12, 21–27^. Notably, PCM1^-/-^ mice exhibit developmental defects and psychomotor and cognitive abnormalities, mirroring those observed in ciliopathies and schizophrenia ^8, 23, 28^. These findings highlight CS as critical regulators of development and differentiation.

Although defined by PCM1, CS are highly heterogenous across cell types and states ^18, 19, 29^. In multiciliated epithelial cells, they appear as granules ranging from 1-6 µm, while in myotubes, they form a matrix surrounding the nuclear envelope, and in oocytes, they are part of a liquid-like spindle domain ^27, 30–32^. CS also undergo dynamic remodeling in response to diverse stimuli. For example, they disassemble during mitosis and reassemble upon mitotic exit ^18, 33^. Their composition and size change during primary cilium assembly and under cellular stress ^26, 34–36^. Additionally, individual CS granules within the same cell differ in size and composition ^13, 37^. This observed intercellular and intracellular reflects their plastic and adaptive nature.

Despite their broad significance, the mechanisms underlying CS assembly, and homeostasis remain poorly understood ^7, 18–20^. Specifically, how PCM1 and clients assemble into functional CS granules with defined size, number, architecture and biochemical properties is unknown. Prior studies provided initial evidence for PCM1’s role in scaffolding CS, and domain analyses showed that its N-terminal half, which contains two self-association domains, can form granules in cells, whereas the C-terminal half cannot^29^. These observations implicated PCM1 multimerization in scaffold assembly, but how this property drives CS formation and how it is regulated in cells is still an open question. Moreover, the intracellular heterogeneity and dynamic remodeling of CS in different cellular states suggest additional layers of regulation, likely through subdomain organization and regulated client recruitment ^27, 29, 38–40^. Yet, the nature and molecular basis of this regulation have not been defined. Dissecting these mechanisms requires tools to monitor CS biogenesis step by step with high spatiotemporal resolution, a challenge given that CS are constitutively present in cells (except during mitosis) and are small relative to the resolution of conventional microscopy.

To overcome these challenges, we developed cellular and *in vitro* CS biogenesis assays that allow for spatiotemporal analysis of the compositional, morphological and dynamic properties of CS granules throughout their biogenesis and maintenance. Using these assays, we show that CS assemble through a hierarchical pathway initiated by PCM1 multimerization followed by client recruitment. We further show that microtubule interactions regulate CS properties, and that perturbing CS granule properties impairs Hedgehog pathway activation. Together, our findings provide mechanistic insight into CS biogenesis and heterogeneity, introduce powerful tools to probe their regulation, and establish a framework for understanding how CS contribute to development and disease.

## Results

### Cellular reconstitution reveals PCM1 multimerization in granule assembly and regulation

An essential first step in elucidating CS assembly mechanisms is characterization of PCM1, as it is the only protein essential for this process. The human *PCM1* gene encodes a 2024 amino-acid (aa) acidic protein that is highly enriched in IDRs (Fig. 1A) ^41^. Its N-terminal 1550 aa region contains eight coiled-coil domains and two self-association domains, while its C-terminal region includes the conserved PCM1-C domain and a LC3-interacting region (LIR) motif (Fig. 1A) ^16, 29, 42–44^. Based on its domain organization, we hypothesized that multimerization of PCM1 drives CS scaffold assembly. To test this in cells, we designed PCM1 deletion mutants that either included or excluded one or both self-association domains. We then examined their ability to form granules in cells, solubility and their association with endogenous PCM1. Specifically, we used GFP-fusions of full-length PCM1 (PCM1) along with deletion mutants PCM1-N (1-1200 aa), PCM1-NS (1-700 aa), PCM1-M (700-1200 aa) and PCM1-C (1200-2024 aa) (Fig. 1B).

**Figure 1.**
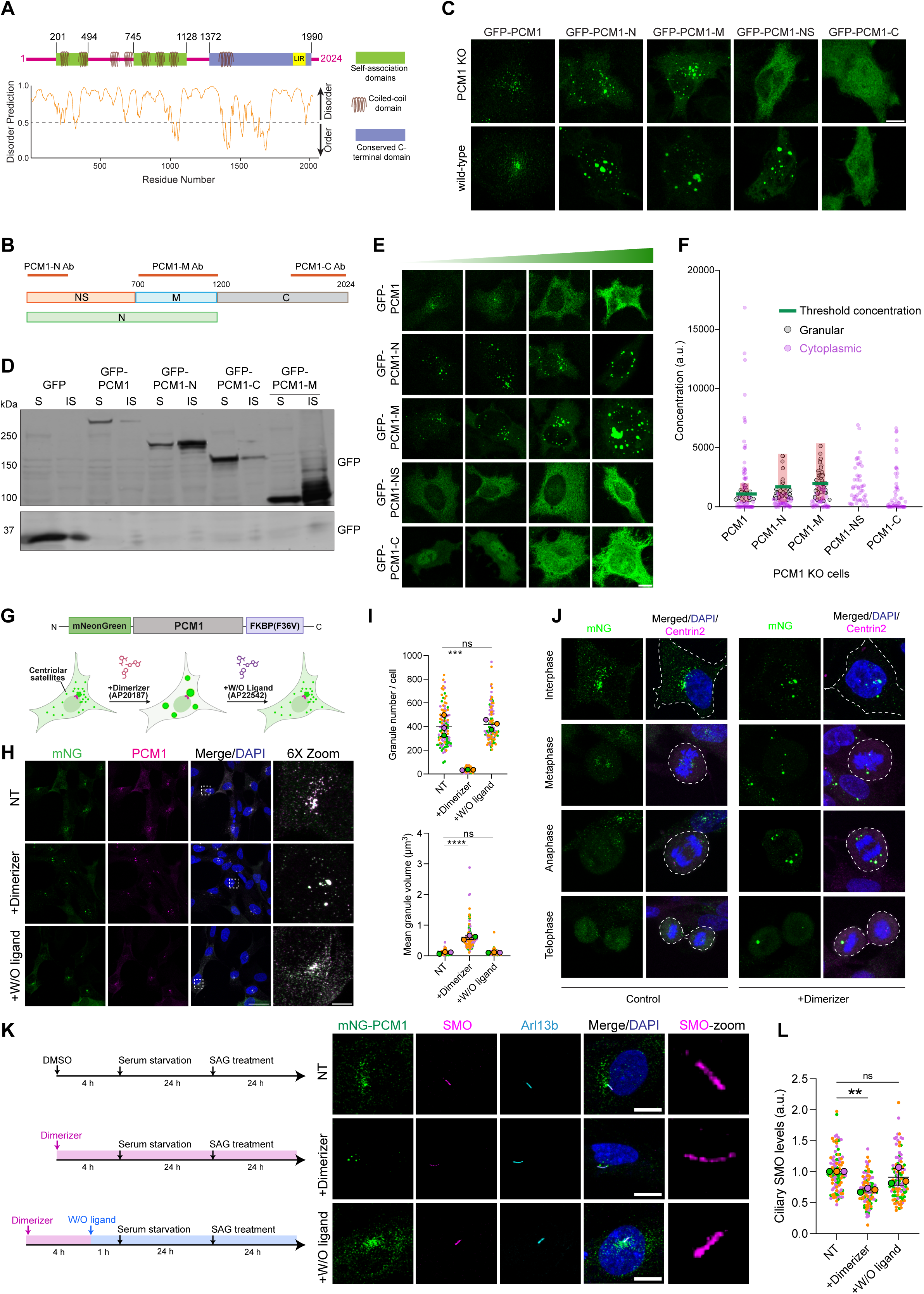
PCM1 is a modular protein with domains important for granule formation and regulation of granule properties. **(A)** Schematic representation of the domain architecture of human PCM1. The *PCM1* gene encodes a 2024 amino acid (aa) acidic protein with a molecular weight of 228 kDa and an isoelectric point (pI) of 4.99. It contains 8 coiled-coil domains, each represented by gray coils located at CC1: 270–293 aa, CC2: 493–513 aa, CC3: 652– 682 aa, CC4: 732–769 aa, CC5: 826–853 aa, CC6: 989–1016 aa, CC7: 1065–1085 aa, and CC8: 1517–1537 aa. It is highly enriched in intrinsically disordered regions (IDRs) predicted by PONDR (predictor of naturally disordered regions) with the VL3 model. Scores above 0.5 indicate disordered regions, whereas scores below 0.5 are ordered. PCM1 contains two self-association domains (201–494 aa and 745–1128 aa, in green), a conserved C-terminal domain (1372–1990 aa, in blue), and an LC3-interacting region (LIR) domain (1953–1958, in yellow). **(B)** Schematic representation of PCM1 truncation constructs designed based on domain analysis and the presence or absence of self-association domains. PCM1-NS contains the first self-association domain, PCM1-M contains the second, and PCM1-N includes both, whereas PCM1-C lacks them. Three antibodies targeting PCM1 at different regions are shown in red (PCM1-N-Ab targets 1-254 aa, PCM1-M-Ab targets 700-1200 aa, PCM1-C-Ab targets 1664-2024 aa). **(C)** PCM1 truncation mutants and their subcellular localization. Representative confocal images show HeLa-WT and HeLa::PCM1 KO cells transfected with the indicated constructs. Cells were stained for anti-GFP. Similar results were obtained from three biological replicates. Scale bar: 10 μm. **(D)** Solubility analysis of GFP-PCM1 and truncation mutants. Immunoblot analysis of lysates from HEK293T cells transfected with GFP, GFP-PCM1, GFP-PCM1-N, GFP-PCM1-C, and GFP-PCM1-M plasmids for 48 hours. Cells were lysed with pulldown buffer and separated as the soluble fraction (S). Remaining insoluble fraction was further lysed with 2X sample buffer. Membranes were stained with anti-GFP-Ab. S: Soluble, IS: Insoluble. **(E)** Localization of PCM1 and its truncation mutants at increasing concentrations in HeLa::PCM1 KO cells. These cells are transfected with GFP fusions of PCM1, PCM1-N, PCM1-M, PCM1-NS and PCM1-C for 48 hours, fixed and stained with anti-GFP. The images indicate the cells with increasing expression levels of PCM1 and its truncations, indicated by a green triangle at the top. Scale bar: 10 μm. **(F)** Quantification of the threshold concentrations at which granules formed for PCM1 and its truncations from (E). For each transfected cell, intensity of GFP signal and total volume and intensity of granules and cytoplasm were measured using Imaris Software. Threshold concentration denotes the mean value of granular concentration values. Results are from three biological replicates, min 48 cells in total. a.u.: arbitrary unit. **(G)** Schematic representation and validation of the chemically inducible CS dimerization system. mNeonGreen-PCM1 is fused to the FKBP(F36V) mutant, which dimerizes upon the addition of AP20187 (homodimerizer) and dimerization is reversed with AP22542 (wash-out ligand) treatment. **(H)** Chemically inducible homodimerization of PCM1 alters CS granule properties. The images represent CS granules under three conditions in RPE1::PCM1 KO cells stably expressing mNG-PCM1-FKBP(F36V): no treatment (NT), with the dimerizer, and with the wash-out (W/O) ligand after dimerization. Cells were treated with the dimerizer for 4 hours and then either fixed directly or treated with the wash-out ligand for 1 hour before fixation. Cells were stained with antibodies for mNeonGreen and PCM1. Scale bars: 30 μm (main panels), 5 μm (insets) **(I)** Quantification of CS granule properties from (H). Granule number per cell and mean granule volume were quantified using Imaris Software. Results are from three biological replicates, green, magenta and orange colored dots representing different replicates. n=35 cells for each replicate. **(J)** Characterization of CS granules following inducible dimerization at various stages of mitosis. Cells were fixed and stained for mNeonGreen and Centrin 2. Representative images show CS granules in control and dimerizer-treated cells at interphase, metaphase, anaphase, and telophase. Scale bar: 10 μm. **(K)** The effect of disrupting CS granule properties on ciliary signaling. The images represent CS granules under three conditions in RPE1::PCM1 KO cells stably expressing mNG-PCM1-FKBP(F36V): no treatment (NT), with the dimerizer, and with the wash-out (W/O) ligand after dimerization. Ciliation was induced by serum starvation for 24 hours. The Sonic Hedgehog pathway was activated by treating cells with 50 nM of the Smoothened agonist SAG for 24 hours. Cells were fixed and stained for mNeonGreen, SMO, and Arl13b. Scale bars: 20 μm (main panels), 5 μm (insets) **(L)** Quantification of Smo recruitment to the cilium as shown in (K). Smo intensity at the cilium was calculated using ImageJ. Only mNG-PCM1-positive cells were selected for quantification. The data represent the mean from three biological replicates, green, magenta and orange colored dots representing different replicates. ns: not significant.

We first performed cellular reconstitution experiments to identify the PCM1 regions required for granule assembly in Hela::PCM1 KO cells (Fig. 1C). GFP-PCM1 formed granules similar to those of endogenous PCM1, clustering around the centrosomes and distributing to a lesser extent throughout the cytoplasm (Fig. 1C). GFP-PCM1-N and GFP-PCM1-M also induced granule formation (Fig. 1C). However, these granules were much larger and more variable in size than those formed by PCM1. Importantly, despite having a self-association domain, GFP-PCM1-NS did not induce granule formation in these cells (Fig. 1C). Similarly, GFP-PCM1-C did not form granules (Fig. 1C). Solubility analysis further showed that PCM1-M and PCM1-N are more insoluble compared to PCM1 and PCM1-C (Fig. 1D), consistent with their tendency to form larger, irregular granules. To assess the influence of endogenous PCM1 on granule formation, we performed similar experiments in wild-type HeLa cells. All mutants showed consistent behavior except for PCM1-NS, which formed granules, likely through multimerization with endogenous PCM1 (Fig. 1C). Supporting this, granules formed by PCM1-N, PCM1-M, and PCM1-NS in wild-type cells recruited endogenous PCM1, confirming their interaction at the CS granules, whereas PCM1-C did not (Fig. S1A).

We next examined the concentration-dependent behavior of PCM1 and its deletion mutants by quantifying the threshold concentrations at which granules formed and how their properties changed with increasing expression levels. To achieve this, we took advantage of variability in expression levels in transiently transfected PCM1 KO cells and measured relative protein concentrations in granules and cytoplasm using fluorescence intensity per volume after 3D reconstruction (Fig. 1E, 1F, S1B) ^45^. Because absolute intracellular concentrations cannot be inferred from fluorescence, we interpreted these measurements as relative values (Fig. S1B). We found that GFP-PCM1 formed granules once it reached a threshold concentration, and these granules dissolved when expression increased further, indicating regulation of CS size and number (Fig. 1E, 1F). In contrast, once GFP-PCM1-N and GFP-PCM1-M granules formed, they did not dissolve at higher concentrations but instead became larger and brighter (Fig. 1E, 1F). In wild-type cells, we were also able to estimate a threshold expression level for GFP-PCM1-NS. The relative thresholds of full-length PCM1 and the mutants differed from those in KO cells, suggesting varying contributions of multimerization with endogenous PCM1 in facilitating granule formation (Fig. S1C, S1D).

Taken together, these findings suggest that PCM1 is a modular protein with PCM1-M being the primary multimerization region required for CS granule formation. Meanwhile, PCM1-C modulates granule properties such as size, number, and solubility, but is not essential for assembly.

### Manipulation of PCM1 multimerization deregulates CS properties and leads to signaling defects

Having established that PCM1 is modular, with PCM1-M driving multimerization and PCM1-C modulating granule properties, we next asked whether altering PCM1’s multimerization state could change CS granule properties and lead to aberrant assembly or disassembly. If PCM1 multimerization is indeed essential for scaffold formation, then perturbing it should alter granule size, number, or stability. To test this, we developed a tool for inducible manipulation of PCM1 multimerization.

To manipulate PCM1 multimerization in cells, we adapted a chemically inducible dimerization system using a modified FKBP protein (FKBP-F36V mutant), which dimerizes upon addition of AP20187 (dimerizer) and whose effect can be reversed with AP22542 (washout ligand) (Fig. 1G) ^46, 47^. FKBP12-F36V homodimerizes upon binding to the dimerizer, which can be rapidly (within several min) reversed with washout ligand. To adapt this system for CS, we generated RPE1::PCM1 KO cells stably expressing mNeonGreen (mNG)-PCM1 fused at its C-terminus to FKBP-F36V (Fig. 1H). We then compared CS granule number and size in these cells before and after dimerization, and following dimer dissociation. Dimerizer treatment reduced granule number and increased their size, which were reversed by washout ligand, confirming that regulated dimerization can control CS size and number (Fig. 1I). Live imaging of mNG-PCM1-FKBP after inducing dimerization and its reversal revealed the spatiotemporal dynamics of these changes (Fig. S2A). These findings show that the inducible dimerization system provides a robust and reversible tool to manipulate CS properties in living cells.

We next examined how inducible dimerization affects the dynamic behavior of CS during mitosis, when CS disassemble in a DYRK-kinase–dependent manner ^40^. As expected, mNG-PCM1-FKBP dissolved during mitosis in untreated cells (Fig. 1J). However, in cells treated with dimerizer, mNG-PCM1-FKBP granules persisted throughout mitosis (Fig. 1J). Some mitotic cells showed granules partitioned into both daughter cells, while others retained them in only one daughter cell (Fig. 1J). A similar pattern of mitotic behavior was observed by time-lapse imaging (Fig. S2B). Thus, induced PCM1 dimerization prevented CS disassembly during mitosis and caused unequal distribution between daughter cells.

Because this inducible system altered CS size and number, it provided a way to directly examine the functional consequences of disrupting CS regulation. In contrast to previous studies that that examined CS function through PCM1 loss, this approach allowed us to probe CS function while maintaining the PCM1 ^8, 10, 11^. Given CS’s well-characterized functions at the cilium in these earlier loss-of-function studies, we asked whether induced dimerization would affect CS functions in cilium assembly and ciliary signaling. Ciliogenesis and cilium length were comparable between control cells and those treated with dimerizer or washout ligand (Fig. S2C-E). However, cilia formed in after dimerizer treatment did not respond efficiently to the Hedgehog agonist SAG, as shown by a significant decrease in ciliary SMO concentration (Fig. 1K, 1L). Importantly, this phenotype was reversed by dissociating the mNG-PCM1-FKBP dimers with the washout ligand (Fig. 1K, 1L). These results demonstrate that manipulating PCM1 multimerization disrupts the formation of functional cilia and link CS regulation to ciliary signaling.

### Development and validation of the *de novo* CS biogenesis assay

The mechanisms underlying CS assembly and its regulation are poorly understood, limiting our ability to explain how CS heterogeneity is established and remodeled in different contexts to support diverse cellular functions. To address this gap, we developed a *de novo* CS biogenesis assay that enables the spatiotemporal analysis of CS granule assembly, including their compositional, morphological and dynamic properties.

Our approach was based on inducible stable expression of GFP-tagged PCM1 and granule-forming deletion mutants (PCM1-N and PCM1-M) in RPE1::PCM1 KO cells. We did not include PCM1-C and PCM1-NS as they failed to form granules in reconstitution experiments (Fig. 1C). To validate our approach, we first assessed whether PCM1 expression induces CS granule assembly. Immunofluorescence analysis after 16 h of induction confirmed the formation of PCM1, PCM1-N, and PCM1-M granules (Fig. S3A). To validate temporal expression of PCM1 in the stable lines, we performed immunoblotting before induction and at 1, 2, 4, 8, and 16 h, which showed a gradual increase in PCM1 levels starting as early as 1 h (Fig. S3B).

To evaluate the physiological relevance of this system, we examined the extent to which the *de novo* formed granules by GFP-tagged PCM1, PCM1-N and PCM1-M were functional. To this end, we performed phenotypic rescue experiments for CS-linked ciliary functions, including cilium assembly and length, recruitment of IFT88 to the basal body and cilium, and Hedgehog pathway activation after SAG stimulation ^8, 10, 11^.

Induced expression of PCM1 rescued defects in ciliogenesis efficiency, cilium length, and ciliary IFT88 and SMO recruitment in PCM1 KO cells, thereby validating that the de novo assay recapitulates physiological CS functions (Fig. S3C-I). Expression of PCM1-N also rescued ciliogenesis but only partially restored cilium length and IFT88/SMO recruitment, indicating that PCM1-N granules are partially functional (Fig. S3C-I). In contrast, GFP-PCM1-M expression failed to rescue any of the ciliary phenotypes, indicating that granule formation alone is not sufficient to support CS-dependent ciliary functions (Fig. S3C-I).

Taken together, these results validate the *de novo* biogenesis assay as a tool to examine CS assembly and function. While GFP-PCM1 fully restored the defects of PCM1 KO cells, PCM1-N and PCM1-M provided partially functional and dysfunctional granules, respectively. These lines therefore allow us to dissect how different PCM1 domains contribute to CS biogenesis.

### Spatiotemporal dynamics of PCM1 scaffold formation during CS biogenesis

CS can assemble either by random self-organization, where scaffold and clients are stochastically recruited from the cytoplasm, or by an ordered pathway, where a scaffold forms first and then recruits clients in a regulated manner. The tightly controlled CS granule properties and heterogeneity observed within and across cell types are more consistent with the latter model ^7, 13, 29^. To test this, we examined scaffold assembly during CS biogenesis. We used the de novo assay to define the spatiotemporal dynamics of PCM1 scaffold formation, and complemented it with mitotic remodeling, a physiological context in which CS undergo disassembly and reassembly during the cell cycle ^33^. For quantitative analysis, we developed an image-analysis pipeline that measured CS number, volume, intensity and distance from the centrosome (Fig. S3J).

Using the *de novo* CS biogenesis assay, we first performed fixed 3D confocal imaging to quantify PCM1 scaffold properties at single-cell resolution before and after doxycycline (dox) induction (Fig. 2A, 2B). Cells were imaged at 10, 20 and 30 minutes, and 1, 2, 4, 6 and 16 hours post-induction. Staining with the centriole marker centrin showed that GFP-PCM1, GFP-PCM1-N, and GFP-PCM1-M granules initially appeared at or near the centrosome. Over time, additional granules of variable size formed in the cytoplasm and gradually clustered around centrosomes, as reflected by decreasing average distance to centrioles. Notably, GFP-PCM1 granules were smaller, more uniform, and more numerous, whereas PCM1-N and PCM1-M formed fewer, larger, and more heterogeneous granules (Fig. 2A, 2B).

**Figure 2.**
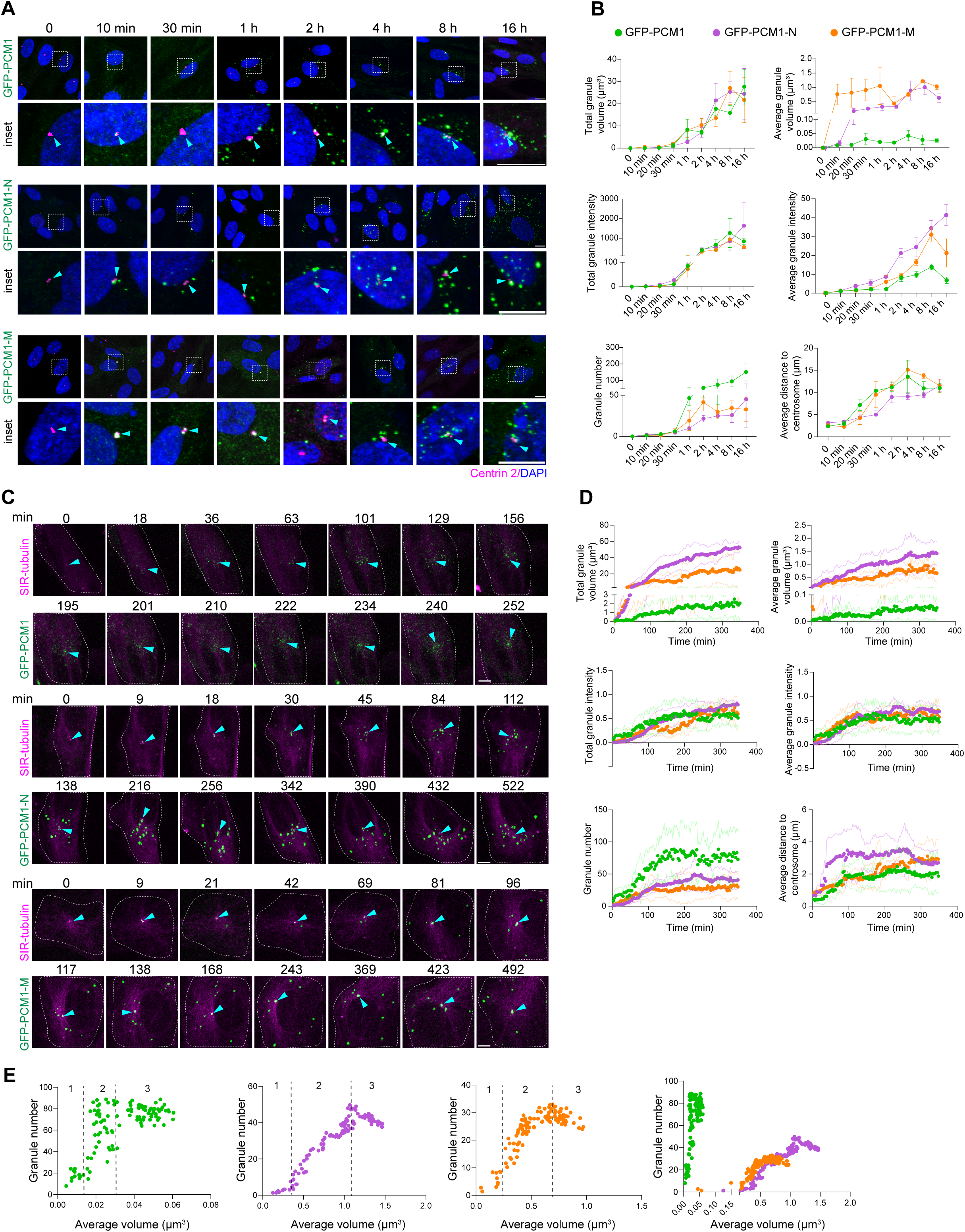
Spatiotemporal dynamics of PCM1 scaffold formation during CS biogenesis. **(A)** Fixed imaging of cells from the *de novo* CS biogenesis assay. Representative confocal images of PCM1 granule formation in RPE1::PCM1 KO cells inducibly expressing GFP-PCM1, GFP-PCM1-N and GFP-PCM1-M. Cells were induced by 1 μM doxycycline for 10 minutes, 30 minutes, 1 hour, 2 hours, 4 hours, 8 hours or 16 hours, fixed and stained with antibodies for GFP and Centrin-2 and DAPI for nucleus. Cyan arrowhead denotes centrosomes at each time point. Scale bars: 10 μm (main panels), 2 μm (insets) **(B)** Quantification of size, fluorescence intensity and number of PCM1 granules from (A). Total volume of granules per cell (μm^3^), average volume of granules per cell (μm^3^), total granule intensity, average granule intensity, granule number, and average distance of granules to centrosome (μm) values were measured using Imaris software. Average values represent total values divided by the number of granules for each cell. Average distance to centrosome values were quantified from the distance between granules and centrosomes marked by centrin 2 for each cell. Results are from three biological replicates, n=48 cells per time point. **(C)** Time-lapse confocal imaging of PCM1 granules from the *de novo* CS biogenesis assay. Images represent still frames from time-lapse confocal images of RPE1::PCM1 KO cells inducibly expressing GFP-PCM1, GFP-PCM1-N, and GFP-PCM1-M. Cells were imaged following the addition of 1 μM doxycycline and SIR-tubulin. The time labeled ’0 min’ marks the initiation of granule nucleation. Centrosomes are indicated by cyan arrowheads at each time point, and cell boundaries are outlined with white dashed lines. Frame times (in minutes) are displayed at the top of each image. Scale bars: 5 μm. **(D)** Quantification of size, fluorescence intensity and number of PCM1 granules from C as described in (B). Average distance to centrosome values were quantified from the distance between granules and SIR-tubulin staining for each cell. Images were taken in 3 minute intervals for 348 minutes. Results are from three biological replicates, data represents the mean of n=5 cells. **(E)** Relationship between average PCM1 granule volume and number during *de novo* CS biogenesis. The average granule volume per cell (μm³) and the number of granules were quantified using Imaris software. Results are from three biological replicates, n=5 cells analyzed.. Assembly progression is categorized into three phases: (1) nucleation, (2) growth, and (3) steady-state.”

To examine spatiotemporal dynamics of assembly, we next performed time-lapse confocal imaging of GFP-PCM1, GFP-PCM1-N, and GFP-PCM1-M assembly dynamics. Quantitative analysis of 3D movies confirmed the fixed-cell observations and revealed three distinct phases of PCM1 assembly (Fig. 2C, 2D). In inducible cells expressing PCM1, an early initiation phase (10–30 min, typically at centrosomes) was followed by a growth phase (30 min–2 h, when granule number, size, and intensity increased), and then by a steady-state phase (after ∼2 h, when granule properties plateaued with modest variation) (Fig. 2C, 2D). PCM1-N and PCM1-M granules also underwent initiation and growth but did not reach a steady state; instead, their size and intensity kept increasing while their numbers decreased, consistent with fusion events ^1^. Plotting granule volume against number confirmed these three phases and showed that PCM1 formed many smaller granules, while PCM1-N and PCM1-M formed fewer, larger ones (Fig. 2E). Together, these analyses reveal a stepwise process of initiation, growth, and steady state during CS biogenesis and highlight that full-length PCM1 and its truncations generate scaffolds with distinct size, number, and intensity profiles.

To validate these findings in a physiological context, we analyzed CS remodeling during mitosis ^33^. Live-cell imaging of GFP-PCM1, GFP-PCM1-N, and GFP-PCM1-M granules with SIR-tubulin labeling revealed a rapid decline in granule number, volume, and intensity at mitotic entry, consistent with disassembly (Fig. S4A, B). As cells progressed through cytokinesis, prominent PCM1 granules reappeared around centrosomes of daughter cells, followed by gradual accumulation of cytoplasmic granules, marking CS reassembly. These dynamics closely paralleled those observed in the *de novo* biogenesis assay.

Together, results from the *de novo* assay and mitotic remodeling experiments revealed distinct stages of PCM1 scaffold assembly and the properties associated with these stages. In both inducible and physiological contexts, PCM1 granules first appear at centrosomes, increase in number during growth phase, consolidate into a steady state, and undergo cell-cycle–regulated disassembly and reassembly.

### Dynamic behavior of PCM1 scaffold during CS biogenesis

We next examined whether PCM1 scaffolds, once formed, maintain dynamic exchange or instead undergo a gradual reduction in dynamics, as has been described for other membrane-less structures such as RNP and pericentrin granules ^45, 48, 49^. Understanding this distinction is important, because the properties of membrane-less organelles often depend on whether their components exchange continuously or are retained in a more static configuration.

To assess the dynamic properties of PCM1 granules, we performed fluorescence recovery after photobleaching (FRAP) analysis. Because the diffraction-limited size of full-length PCM1 granules prevented reliable FRAP measurements, we first examined PCM1-N and PCM1-M. Time-dependent FRAP showed progressively reduced mobile fractions and slower recovery, indicating that these granules gradually lose dynamic exchange (Fig. 3A, 3B). Live imaging of GFP-PCM1-N granules revealed dynamic events among granules, including occasional fusion and splitting, but the majority of interactions were transient “kiss-and-run” contacts where granules briefly touched and then separated again (Fig. 3C). The predominance of kiss-and-run behavior supports that most PCM1-N granules adopt a less dynamic state once assembled.

**Figure 3.**
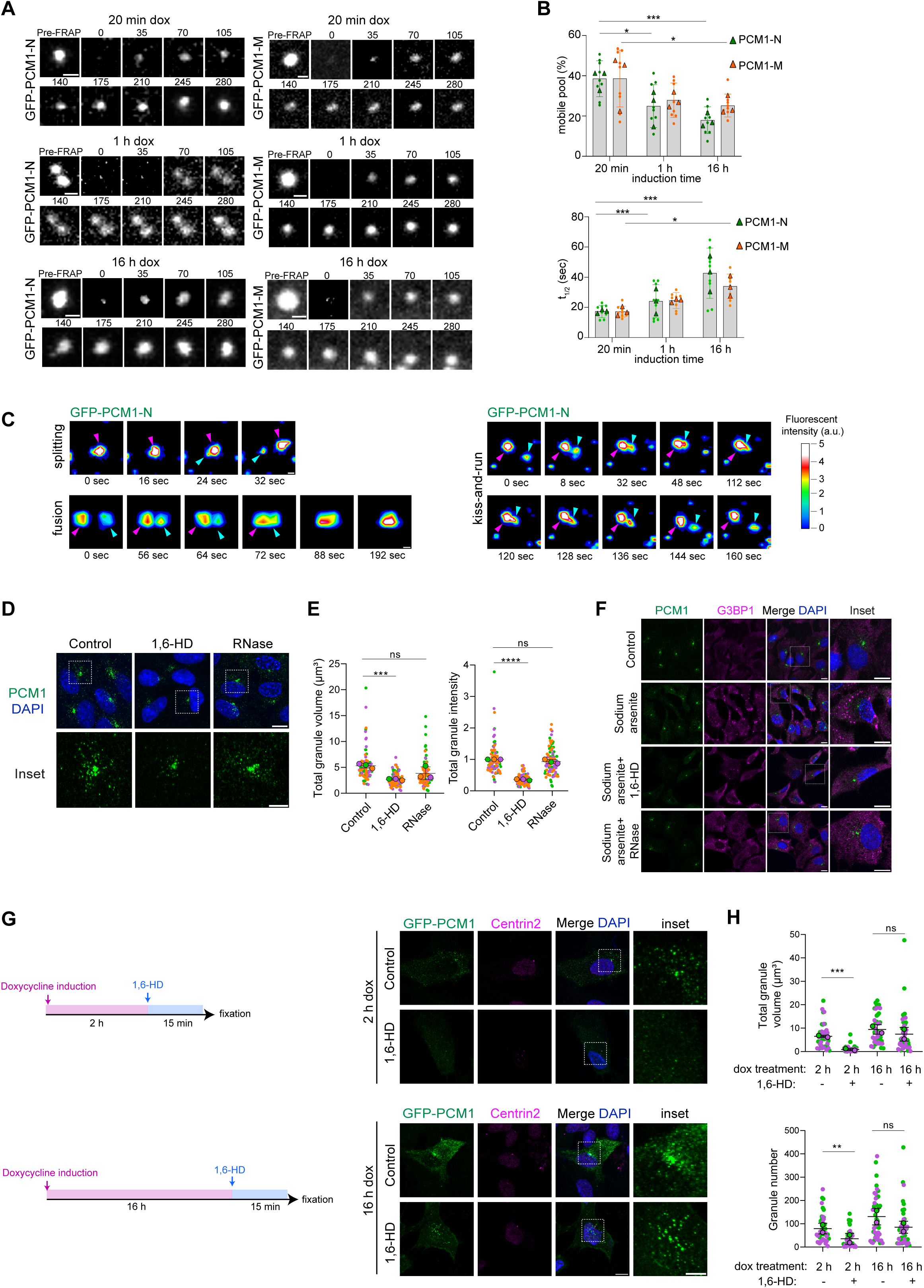
Dynamic properties of CS. **(A)** FRAP analysis on GFP-PCM1-N and GFP-PCM1-M granules formed in a *de novo* biogenesis assay. RPE1::PCM1 KO cells expressing either GFP-PCM1-N or GFP-PCM1-M were induced with doxycycline for 20 minutes, 1 hour, and 16 hours. The specified regions were photobleached and subsequently imaged for 300 seconds post-bleaching. Scale bars: 10 μm (main panels), 1 μm (insets). **(B)** Quantification of mobile pool and half-time of fluorescence recovery from (A). Mobile fractions and half-lives times of fluorescence recovery were calculated based on granule recovery (n = 3 independent FRAP experiments from three biological replicates). Circles represent individual experiments, and rectangles indicate the mean values. **(C)** Representative time-lapse images show the dynamics of fusion, splitting, and kiss-and-run interactions among PCM1 granules. Representative still frames from time-lapse confocal image show there different dynamic event. Granules are marked with magenta and cyan arrowheads. Five different colors show varying levels of fluorescent intensity, ranging from blue (lowest) to white (highest). These results denote one granule as an example for each dynamic behavior and are consistent with findings from three independent experiments. **(D)** Representative confocal images of RPE1 cells treated with different reagents, fixed with methanol and stained with anti-PCM1-C Ab and DAPI for nucleus. Untreated cells serve as the control. Cells were treated with 5% 1,6-hexanediol for 5 minutes, 100 μg/ml RNase A+I for 1 hour. Scale bars: 10 μm (main panels), 5 μm (insets). **(E)** Total volume of granules per cell (μm^3^) and total granule intensity values were measured using Imaris software. Granule intensity was normalized to control values for each replicate. Number of cells for each total intensity value was determined from relative frequency data. Statistical significance was determined by the Student’s *t*-test (paired and two-tailed). Results are from three biological replicates, green, magenta and orange colored dots representing different replicates. n=30 cells for each replicate. \**P*<0.05; \*\**P*<0.01, \*\*\**P*<0.001\*\*\*\**P*<0.0001; ns, not significant. **(F)** Experimental schematic of 1,6-Hexanediol and RNase treatments after stress induction. RPE1 cells were treated with 50 mM sodium arsenate treatment for 1 hour for stress induction, followed by 5% 1,6-Hexanediol treatment for 5 minutes or 100 μg/ml RNase A + I treatment for 1 hour. Cells were then fixed and stained for PCM1 and G3BP1 antibodies marking CS and stress granules respectively. Cells were stained with anti-PCM1-Ab (green), anti-G3BP1 (magenta) and DAPI for nucleus (blue). Scale bars: 10 μm (main panels), 2 μm (insets). **(G)** Representative confocal images of RPE1::PCM1 KO cells inducibly expressing GFP-PCM1 transfected with either control or 1,6-HD treatment after 2 h or 16 h doxycycline induction. After 2 h or 16 h doxycycline induction, cells were treated with 5% 1,6-HD for 5 minute, whereas control cells were fixed after 5 minutes. Cells were fixed and stained for anti-GFP (green), anti-centrin (magenta) and nucleus (blue). Scale bars: 10 μm for images, 5 μm for insets. **(H)** Granule properties, including total volume of granules per cell (μm^3^), average volume of granules per cell (μm^3^), total granule intensity, granule number values were measured from (G) using Imaris software. Average values represent total values divided by the number of values for each cell. Statistical significance was determined by the Student’s *t*-test (paired and two-tailed). Results are from two biological replicates, green and magenta colored dots representing different replicates. *:P<0.05; **:P<0.01, ***:P<0.001****:P<0.0001; ns: not significant.

To extend these findings to full-length PCM1, we examined the behavior of its granules at steady state using chemical and enzymatic perturbations. We treated RPE1 cells with 1,6-hexanediol (1,6-HD), a small aliphatic alcohol that disrupts weak hydrophobic interactions and is widely used to probe the dynamics of membrane-less organelles ^50, 51^. 1,6-HD treatment reduced the total volume and intensity of PCM1 granules, indicating that a small fraction remains sensitive and dynamic (Fig. 3D, 3E).

By contrast, majority of PCM1 granules persisted, showing that most adopt a non-dynamic state once assembled. As a control, 1,6-HD almost completely dissolved stress granules marked by G3BP1, which are known to be highly dynamic, confirming the sensitivity of the assay (Fig. 3F) ^52, 53^. We also treated cells with RNase, as RNA has recently been defined as a CS component and RNA-RNA/RNA-protein interactions often underlie membrane-less organelle assembly ^1, 5, 54^. PCM1 granules remained intact, whereas stress granules dissolved (Fig. 3D-F). Finally, to determine how dynamics change during biogenesis, we compared 1,6-HD sensitivity at early (2 h) and late (16 h) induction time points. PCM1 granules were more sensitive at 2 h, with larger reductions in number and volume than at 16 h, showing that they become progressively less responsive to disruption as they mature (Fig. 3G, 3H).

Together, FRAP, live imaging and chemical perturbation assays show that while a minor fraction of PCM1 assemblies retains dynamic properties, the majority progressively adopt a predominantly non-dynamic state during biogenesis.

### Clients are recruited hierarchically to the PCM1 scaffold

Having established that PCM1 initiates CS scaffold assembly, the next question is how client proteins are incorporated. One possibility is that PCM1 first forms a scaffold that subsequently recruits clients in a stepwise manner, giving rise to functional CS granules with distinct composition. Alternatively, scaffold and clients may co-assemble from the beginning. To distinguish between these possibilities and define the sequence of CS assembly, we examined the timing of client recruitment relative to PCM1 scaffold formation.

Using the *de novo* biogenesis assay with GFP-PCM1, we fixed cells at defined time points after doxycycline induction, stained the clients with specific antibodies and quantified their colocalization with PCM1 (Fig. 4A). For these experiments, we selected nine established CS clients with roles spanning ciliogenesis, centriole duplication, and microtubule organization, several of which are also implicated in human diseases including microcephaly, ciliopathies, and oral–facial–digital syndrome ^7, 11, 17, 19, 37, 55–60^. Of note, our analysis was limited to clients for which antibodies were available that specifically detect their CS pool. Our analysis was restricted to proteins for which antibodies were available that detect their CS pool in addition to their centrosomal pool.

**Figure 4.**
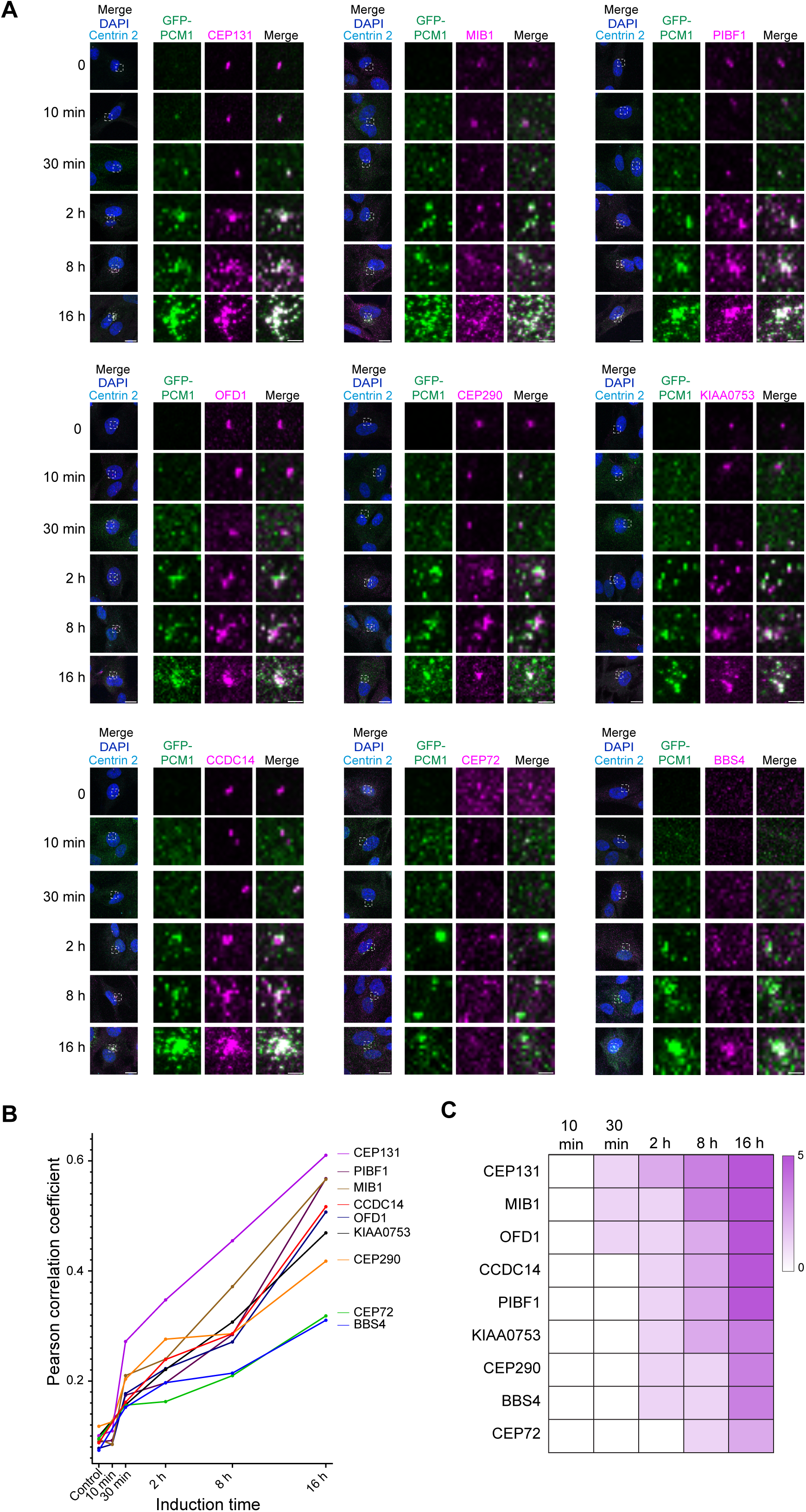
Spatiotemporal analysis of client recruitment during CS biogenesis. **(A)** RPE1::PCM1 KO cells inducibly expressing GFP-PCM1 were fixed before induction and 10 min, 30 min, 2 hours, 8 hours, 16 hours after induction. They were then stained with antibodies for GFP and clients denoted in magenta. Representative confocal images are shown for each client and time point. White dashed boxes highlight zoomed-in regions. Scale bars: 10 μm (main panels), 2 μm (insets) **(B)** Pearson correlation coefficient (PCC) analysis of client-scaffold colocalization. PCCs between PCM1 and client proteins were calculated from 3D images using Imaris Software (intensity/voxel-based). They represent colocalization analysis across 30 cells per condition in two independent experiments. The negative control, represented by the PCC between PCM1 and G3BP1 (0.032), indicates the lower detection limit of our colocalization. The positive control, between PCM1 and PCM1 (0.887), represents the upper limit of our colocalization detection. **(C)** Classification of client recruitment dynamics based on colocalization strength. Recruitment time of each client to scaffold PCM1 was determined by comparing Pearson Correlation Coefficient (PCC) values relative to the 0-minute condition. Colocalization strength was classified as positive if the PCC value at 10 minutes was at least twice that at 0 minutes. Darker magenta color represents up-to 5 times increase in the PCC value compared to 0 minutes. These thresholds were established based on the observed colocalization ranges in our dataset. A 2-fold increase marks the minimum detectable enrichment above baseline, ensuring subtle recruitment events are noted, while a 5-fold increase represents the maximum detected, with 3-fold and 3.5-fold serving as intermediate steps. This data-driven approach captures the full spectrum of recruitment behaviors effectively.

To measure recruitment, we calculated Pearson correlation coefficients (PCC) between PCM1 and each client before induction and at 10 min, 30 min, 2 h, 8 h, and 16 h post-induction (Fig. 4A). Control measurements established the range of detection. PCM1 stained with Alexa 488- and Alexa 568-conjugated secondary antibodies gave a PCC of 0.89 (upper limit), whereas PCM1 and the non-CS protein G3BP1 gave a PCC of 0.03 (lower limit). For each client, we plotted PCC values over time (Fig. 4B) and also compared changes between consecutive time points to infer relative recruitment order (Fig. 4C). Together, these complementary approaches allowed us to follow how clients associate with PCM1 scaffolds during biogenesis.

Our analysis revealed that no clients were detectable at PCM1 granules as early as 10 min, supporting that CS granule formation is driven by PCM1 and does not involve immediate co-assembly with clients (Fig. 4A). By 30 min, several clients including CEP131, CEP290, MIB1, OFD1, and PIBF1 could be detected at the PCM1 granules near centrosomes (Fig 4A-C). Among these, CEP131 consistently showed the strongest correlation, suggesting it is among the earliest recruited clients. By 2 h, all tested clients were detectable at PCM1 granules, although with different PCC levels (Fig. 4B, 4C). Client-PCM1 PCCs increased further at 8 and 16 h, reaching values between 0.3 and 0.6, consistent with intracellular CS compositional heterogeneity (Fig. 4B, 4C). Across all time points, CEP131, MIB1, and PIBF1 showed higher colocalization with PCM1; CCDC14, OFD1, KIAA0753, and CEP290 showed intermediate association; and CEP72 and BBS4 consistently showed the lowest. Importantly, even when clients reached similar PCCs at 30 min or 16 h, their recruitment trajectories differed. For example, CEP131 and PIBF1 both reached high PCCs by 16 h but had distinct initial rates of accumulation (Fig. 4B, 4C). These differences indicate that client recruitment is progressive and client-specific, contributing to intracellular heterogeneity. Because antibody sensitivities vary, it is possible that some apparent differences reflect detection efficiency rather than true recruitment order. Nonetheless, the reproducible variation in PCC profiles across biological replicates supports the conclusion that clients are not incorporated simultaneously but instead associate with PCM1 scaffolds in distinct patterns over time.

To determine how scaffolds formed by different PCM1 domains influence client recruitment, we examined cells expressing GFP-PCM1-N or GFP-PCM1-M at the same time points. Recruitment to PCM1-N largely mirrored that of full-length PCM1 (Fig. S5A– C). Because PCM1-N formed larger granules, additional clients such as ninein, likely below detection threshold with PCM1, were detected at PCM1-N granules. BBS4, which requires the C-terminus of PCM1, was absent from PCM1-N but recruited to full-length PCM1 ^34, 61^. Most clients showed higher PCCs with PCM1-N compared to PCM1, consistent with stronger recruitment to larger granules. By contrast, client recruitment to PCM1-M was highly restricted: only CEP290 localized to PCM1-M granules, and only at later time points (16 h) and in the largest granules, suggesting weak affinity (Fig. S5D). These results indicate that PCM1-M contributes primarily to PCM1 self-association rather than client recruitment, consistent with its inability to restore ciliary defects in PCM1 KO cells (Fig. S3C,D).

### PCM1 and clients are organized into subdomains within CS granules

Internal organization has been described for several membrane-less organelles, including paraspeckles and the pericentriolar material, where core-shell or layered architectures help coordinate protein function ^62–64^. Given the multistep assembly and regulated heterogeneity of CS granules, we hypothesized that they might also contain internal subdomains. To test this, we mapped the spatial arrangement of PCM1 and selected clients using ultrastructure expansion microscopy (U-ExM) and structured illumination microscopy (SIM).

First, we analyzed the distribution of different PCM1 domains using U-ExM (Fig. 5A). Because PCM1 is a large coiled-coil protein, its domains could probe different spatial domains, as seen for centrosome proteins like CEP152 and Plp ^62, 63^. To test this, we used antibodies against the N-terminal (1–254 aa), middle (700–1200 aa), and C-terminal (1664–2024 aa) regions of PCM1, and validated their specificity with deletion mutants (Fig. S6A–D). Fluorescence intensity profiles showed that these regions overlapped but were not identical, suggesting that PCM1 domains occupy partially distinct positions within granules (Fig. 5A).

**Figure 5.**
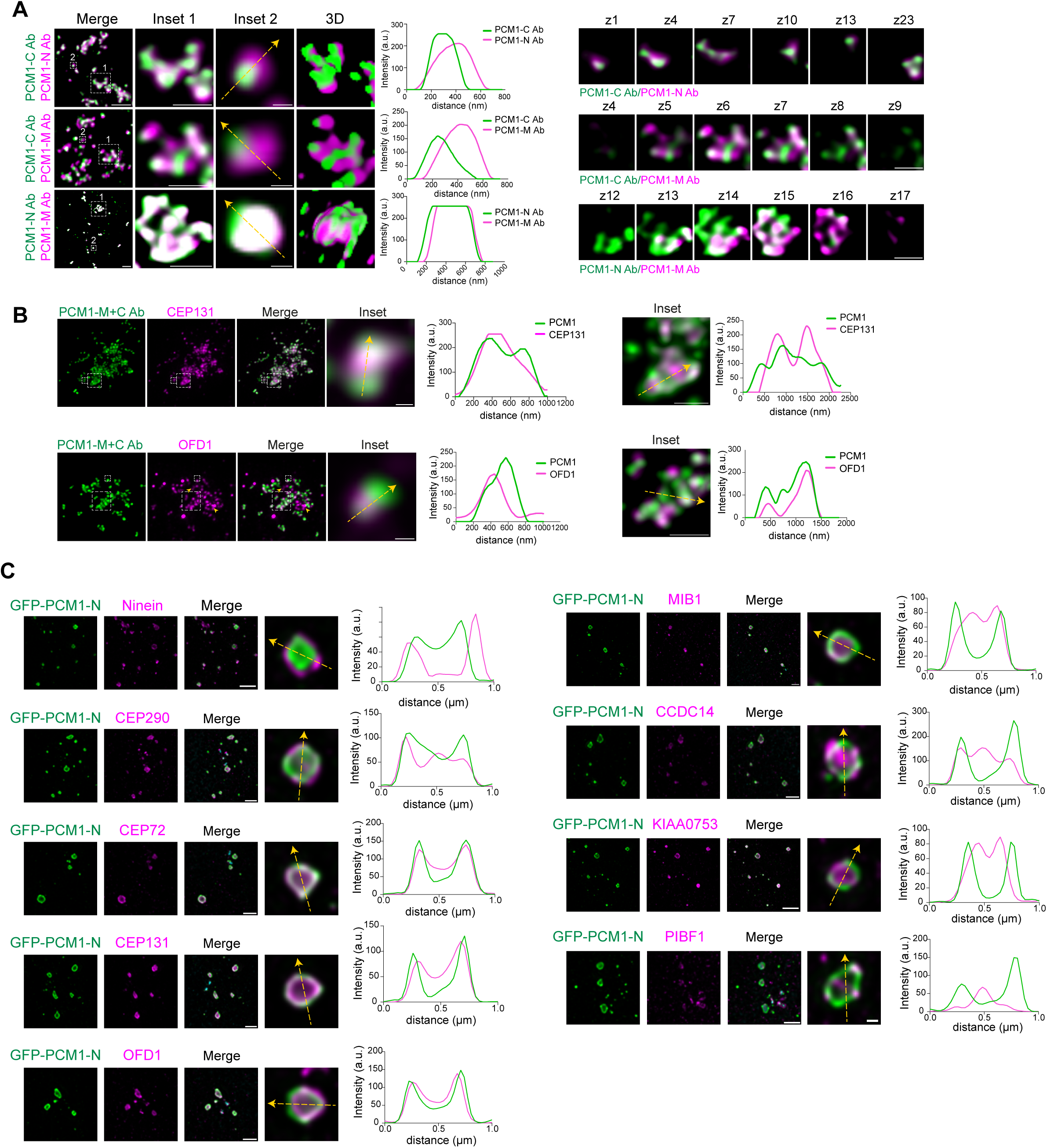
PCM1 and clients are organized into subdomains within CS granules. **(A)** Spatial organization within the PCM1 granules. RPE1 cells were processed for U-ExM and stained for combinations of PCM1-C-Ab, PCM1-M-Ab and PCM1-N-Ab in green or magenta. Scale bars shown on the images are not adjusted for the expansion factor (4.2x). Insets display 4× or 10× magnifications of the cluster or individual CS. Individual Z-stack images depict the spatial organization of PCM1 stained with antibodies for the PCM1-C terminus, PCM1-N terminus, and PCM1-M region. 3D representations are rendered from Leica Software. Scale bars: 2 μm (main panels), 1 μm and 200 nm, respectively (insets). **(B)** Representative confocal images of expanded CS showing the spatial organization of PCM1 and its clients. RPE1 cells were processed for U-ExM and stained for clients (green) anti-CEP131 or anti-OFD1, and anti-PCM1-M-Ab / PCM1-C-Ab mixture (magenta). Scale bars shown on the images are not adjusted for the expansion factor (4.2x). Insets display 4× or 10× magnifications of the cluster or individual CS. Arrows indicate the centrioles. Scale bars: 2 μm (main panels), 1 μm and 200 nm, respectively (insets). **(C)** Spatial organization of PCM1-N and clients within CS granules. Images show high magnification of GFP-PCM1-N granules colocalizing with various client proteins. RPE1::PCM1 KO cells inducibly expressing GFP-PCM1-N were induced for 24 hours and fixed and stained with anti-GFP-Ab (green) and clients (magenta). Cells were imaged using Zeiss Elyra 7 Lattice SIM superresolution microscope. Intensity plot profiles were drawn using ImageJ. Scale bars: 0.2 μm.

We next examined the relationship between PCM1 scaffold and clients. Using U-ExM on RPE1 cells stained with the PCM1 N-terminal and client antibodies, we found that only CEP131 and OFD1 could be reliably detected under these conditions. CEP131 was enriched in a core region surrounded by PCM1, whereas OFD1 localized more peripherally with partial overlap (Fig. 5B). These results indicate that PCM1 and clients are not uniformly distributed but can occupy different zones within granules.

To increase spatial resolution and detect additional clients, we analyzed larger PCM1-N granules, which reach up to 1 μm in diameter, provide more binding sites, and retain partial functionality during cilium formation and signaling (Fig. S3C, S3D). RPE1::PCM1 KO cells expressing GFP-PCM1-N were fixed and stained for the nine clients previously examined in the biogenesis assay (Fig. 5C). Since inducible biogenesis assays revealed that granules increase in size during biogenesis, we imaged structures of different sizes (0.1–1 μm) to determine whether client distribution changes with granule size (Fig. S6E).

GFP-PCM1-N assembled into spheroidal scaffolds, with organization resolvable in granules larger than 0.4 μm (Fig. 5C, S6E, S6F). In smaller granules, differences between PCM1-N and clients were visible but precise positioning was less clear. In larger granules, intensity profiles suggested three broad categories of client localization relative to PCM1-N: core proteins (CEP131, KIAA0753, PIBF1), enclosed within a continuous PCM1 shell; shell proteins (CEP290, CEP72), overlapping with PCM1 at the periphery, with ninein localized externally; and patch proteins (e.g., CCDC14, PIBF1), present as puncta within the scaffold. Some proteins, including OFD1 and MIB1, localized to both the core and shell (Fig. 5C, S6E).

Taken together, these analyses suggest that CS granules exhibit a core–shell– patch arrangement in which PCM1 and clients occupy distinct subdomains. While antibody availability and resolution limit the breadth of this analysis, the reproducible distribution patterns indicate that CS granules are not homogeneous. This substructuring may provide a basis for compartmentalization and regulation of client proteins, contributing to the heterogeneity and adaptability of CS.

### CS subdomains differ in client interaction dynamics and composition

The identification of distinct CS subdomains and the predominantly non-dynamic nature of the PCM1 scaffold raised the question of whether scaffold–client associations vary in their dynamics and whether CS composition remodels in different cellular states. Addressing this is important because such regulation could explain the heterogeneity of CS granules and their ability to adapt to diverse cellular functions.

To probe interaction dynamics, we perturbed weak hydrophobic interactions among scaffold and clients with 1,6-HD. Co-immunoprecipitation showed that the association of CEP131, MIB1, and KIAA0753 with PCM1 decreased after treatment, whereas the PCM1–CEP290 interaction was largely unaffected (Fig. 6A). This indicates that some clients interact with PCM1 through more dynamic associations, while others associate less dynamically.

**Figure 6.**
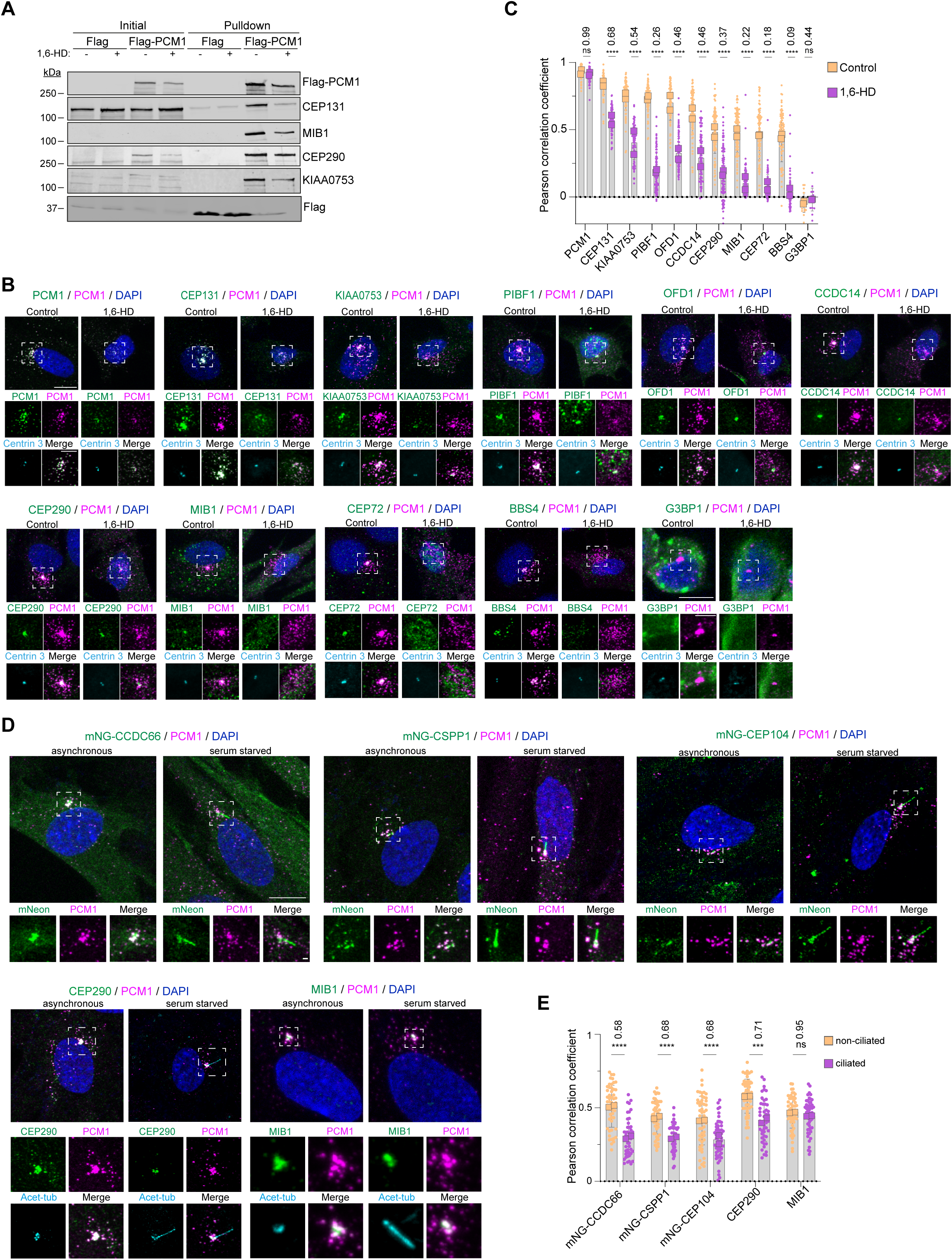
CS subdomains and granules have distinct compositions and biophysical properties. **(A)** Control and 1,6-hexanediol-treated HEK293T::PCM1 KO cells transfected with either FLAG or FLAG-PCM1 were lysed, and proteins were pulled down by Flag beads. The initial sample (initial) and pulled down proteins were run on an SDS gel and immunoblotted with antibodies against Flag-PCM1 and clients. **(B)** RPE1 cells were treated with 5% 1,6-hexanediol for 10 min, fixed with methanol, and stained with antibodies against the indicated client proteins (green), PCM1 (magenta), centrin 3 (to mark the centrosome), and DAPI (to visualize DNA). A dual-secondary antibody labeling strategy was used to confirm PCM1 detection in both channels (top left panel). Cells in the bottom right panel were additionally treated with 50 mM sodium arsenate for 1 h to induce stress granules before 1,6-hexanediol treatment. Confocal microscopy was used for imaging. White boxes indicate zoomed-in regions. Scale bars: 10 μm (main panels), 5 μm (insets). **(C)** Pearson correlation coefficient quantifying colocalization between PCM1 and client proteins. Small circles represent individual cells, while large circles indicate the average per experiment. Fold change is calculated by comparing the mean value of 1,6-hexanediol -treated samples to that of the control. Results are from two biological replicates, n = 40 cells per condition. Statistical significance was determined using a t-test comparing treatment and control for each colocalization pair (ns > 0.05, ****P < 0.0001). **(D)** Asynchronous and serum starved RPE1::mNeonGreen-CCDC66, RPE1::mNeonGreen-CSPP1 and RPE1::mNeonGreen-CEP104 cells were fixed with methanol, and stained with antibodies against mNeonGreen (green) and PCM1 (magenta). Additionally, asynchronous and serum starved RPE1 cells were fixed with methanol, and stained with antibodies against the indicated client proteins (green), PCM1 (magenta) and acetylated tubulin (cyan) (to mark the primary cilia). White boxes indicate zoomed-in regions. Scale bars: 10 μm (main panels), 1 μm (insets). **(E)** Pearson correlation coefficients for quantifying colocalization between PCM1 and client proteins. Fold change is calculated by comparing the mean value of serum starved cells to the mean of asynchronous cells. Results are from two biological replicates, n = 30 cells per condition. Statistical significance was determined using a t-test comparing treatment and control for each colocalization pair (ns > 0.05, **P < 0.01, ****P < 0.0001).

We then asked whether such differences contribute to compositional heterogeneity within cells ^13, 37^. Using PCC analysis, we quantified client–PCM1 colocalization before and after 1,6-HD treatment. The association of all tested clients decreased but to varying degrees (Fig. 6B, 6C). CEP131, KIAA0753, OFD1, and CCDC14 showed moderate reductions (30–55%), PIBF1, CEP290, and MIB1 were more strongly affected (60–80%), and CEP72 and BBS4 showed the largest decreases (80–99%). These graded responses likely reflect differences in interaction dynamics.

Notably, CEP72 and BBS4 were also the last clients to be recruited during CS biogenesis (Fig. 4), consistent with more transient associations. In contrast, RNase treatment had little effect on client–PCM1 association, except for CEP131, which may partially depend on RNA for recruitment (Fig. S7A, S7B).

Finally, we tested whether CS composition remodels in response to cellular context. Comparing unciliated and ciliated RPE1 cells, we observed reduced PCM1 association of ciliogenesis regulators, including positive regulators CEP290, CCDC66, CSPP1, and CEP104, whereas MIB1 remained unchanged (Fig. 6D, 6E) ^11, 35, 65, 66^. These client-specific differences suggest that CS composition is not static but can remodel depending on cellular state, selectively reducing the association of positive ciliogenesis regulators when cilia are established. Together, these results suggest that scaffold–client associations differ in their dynamics and that CS are heterogeneous organelles that adapt to cellular requirements.

### *In vitro* reconstitution reveals scaffold formation, client recruitment and microtubule interaction as intrinsic PCM1 properties

CS functions rely on their microtubule-based positioning and transport and selective recruitment of clients. Our cellular assays demonstrated that PCM1 multimerization is required for scaffold assembly and that this scaffold recruits clients during CS biogenesis. What remains unknown is whether these properties are intrinsic to PCM1 or require additional cellular factors. To directly address this, we performed *in vitro* reconstitution experiments with purified PCM1, clients, and microtubules.

We expressed and purified His-MBP-mNeonGreen-PCM1 (referred to as mNG-PCM1) from insect cells and verified the presence of full-length protein by immunoblotting and immunostaining with antibodies specific to its N-and C-terminal regions (Fig. S8A, S8B). These antibodies bound to distinct subdomains within the precipitated PCM1 protein, consistent with its modular organization (Fig. S8B). Mass spectrometry further confirmed that mNG-PCM1 was the major protein present, with no enrichment of centrosome or microtubule regulators (Fig. S8C, Table S1).

We next tested whether purified PCM1 can form granules under buffer conditions mimicking the cytoplasm. At 1 μM, mNG-PCM1 rapidly assembled into spherical granules in a low-salt buffer containing 150 mM NaCl (Fig. 7A). This protein concentration approximates intracellular PCM1 levels (∼700–900 nM in mammalian cells) and therefore mimics the cellular environment ^67–69^. Notably, the N-terminal mNG tag appeared in a spheroidal arrangement, particularly evident in larger granules, suggesting spatial organization within PCM1 granules (Fig. 7A inset). Increasing protein concentrations from 1 to 4 μM increased granule number but not size, indicating size regulation (Fig. 7B, 7C). As a negative control, mNG alone did not form granules (Fig. S8D).

**Figure 7.**
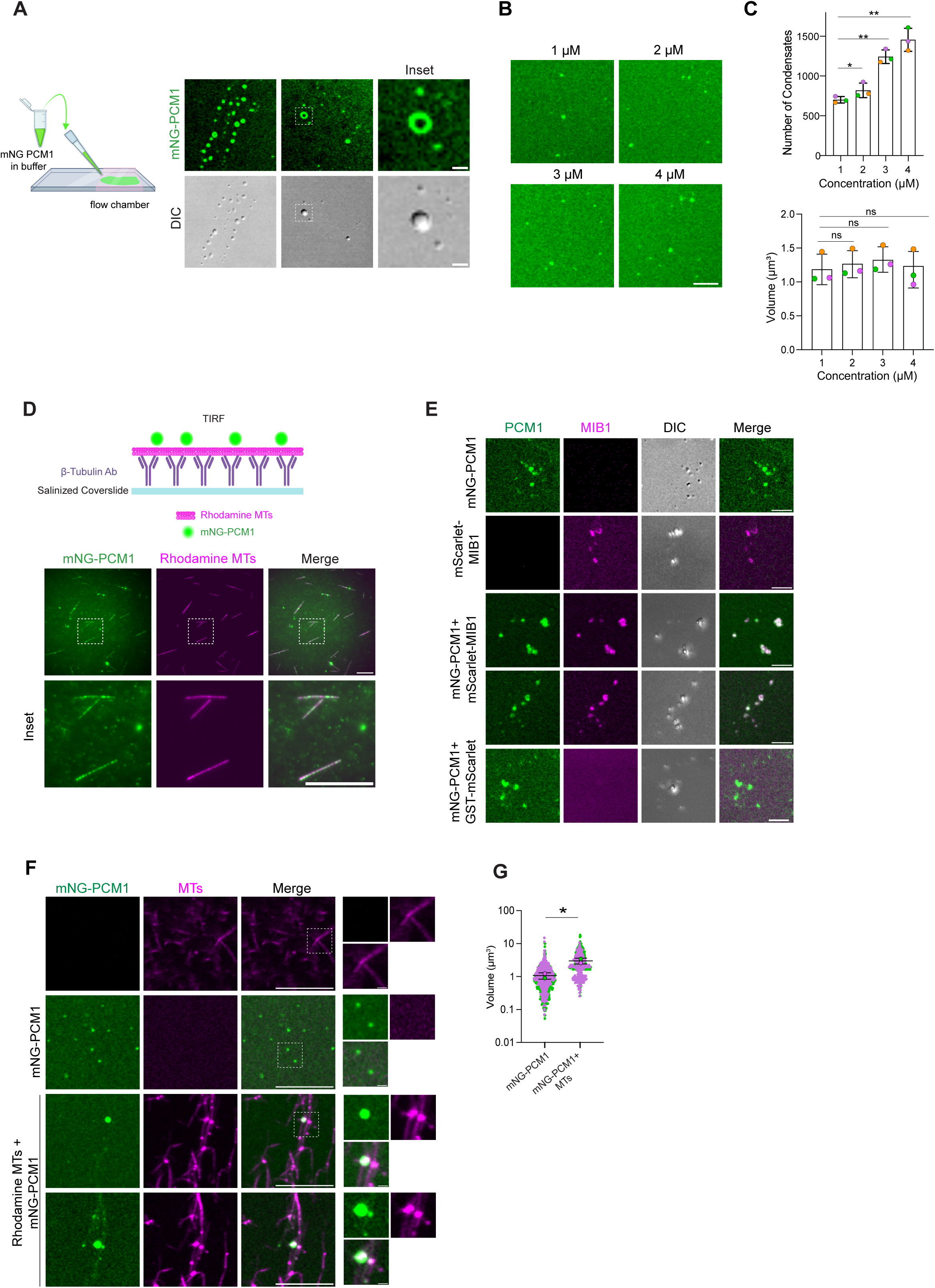
PCM1 exhibits intrinsic properties for granule formation, client sequestration, and microtubule binding *in vitro*. **(A)** Schematic depicts the experimental flow. mNeonGreen-PCM1 condensates in a phase separation buffer at 1 μM, flown in chambers and imaged using a confocal microscope. Scale bar, 2 μm for field of view. **(B)** mNeonGreen-PCM1 condensates in phase separation buffer at concentrations 1, 2, 3, 4 μM. **(C)** Quantification of the number of condensates and volumes of each granule (μm^3^) in the phase separation buffer from D. Results are from three biological replicates. **(D)** mNeonGreen-PCM1, mScarlet-MIB1 and GST-mScarlet were set individually and combined in phase separation buffer, flown in chambers and imaged. Scale bar, 5 μm. **(E)** Schematic depicts the experimental flow. Taxol stabilized microtubules polymerized from purified bovine tubulin labeled with Alexa Fluor 568 were attached on salinized and tubulin antibody coated cover slide surface. mNeonGreen-PCM1 was flown in a chamber and imaged with TIRF microscopy. Scale bar, 10 μm (main panels); 2 μm (insets). **(F)** mNeonGreen-PCM1 and taxol-stabilized microtubules, polymerized from purified bovine tubulin and labeled with Alexa Fluor 568, were prepared in BRB80 buffer both individually and in combination. These were introduced into flow chambers and imaged. Scale bars: 10 μm (main panels), 1 μm (insets). **(G)** Volume of mNeonGreen-PCM1 condensates plotted for each condition, with the y-axis on a log10 scale for the mNeonGreen-PCM1 only and the additive microtubules condition. Results are from two biological replicates, green and magenta colored dots representing different replicates.

We then investigated whether PCM1 granules *in vitro* recapitulate key cellular properties of CS. We first examined their ability to recruit clients. Because of its established role in CS homeostasis and function and its characterization in our biogenesis assays, we chose MIB1 as a candidate client ^11, 70^ (Fig. S8E). Purified His-MBP–mScarlet–MIB1, added at ∼100 nM (reflecting cellular levels) ^67–69^, could form granules on its own, but it was also selectively incorporated into pre-formed PCM1 granules (Fig. 7D). In contrast, GST–mScarlet was not incorporated (Fig. 7D). These results show that PCM1 scaffold can directly and selectively recruit client proteins *in vitro*, with MIB1 serving as an example of selective recruitment.

Next, we tested for microtubule interaction, given that CS are transported along microtubules and this is required for their functions ^9, 71^. TIRF microscopy showed that mNG-PCM1 granules bound directly to taxol-stabilized, rhodamine-labeled microtubules immobilized on coverslips (Fig. 7E). When mNG-PCM1 was incubated with rhodamine-labeled microtubules in solution, microtubules reorganized into denser bundles with larger PCM1 granules associated along them (Fig. 7F, 7G, S8F). Notably, these granules were larger than in PCM1-only controls and often exhibited spheroidal organization, suggesting that microtubules might promote scaffold assembly (Fig. 7F, 7G, S8G). Tubulin itself was recruited into PCM1 granules, confirming its role as a CS client and consistent with prior evidence of PCM1–tubulin interactions ^72^. Consistent with microtubules’ role in CS biogenesis, nocodazole treatment in RPE1::PCM1 KO cells expressing GFP-PCM1 compromised scaffold formation. Treated cells displayed fewer and smaller PCM1 granules with lower fluorescence intensity compared to controls, along with redistribution throughout the cytoplasm (Fig. S8G, S8H). These effects were strongest at 2 h, with partial recovery by 16 h.

Together, these assays show that PCM1 intrinsically self-assembles into regulated granules with internal organization, recruits clients such as MIB1 and tubulin, and associates directly with microtubules. In addition, our data indicate that microtubules regulate CS assembly, likely by providing a scaffolding function.

## Discussion

In this study, we combined cellular and *in vitro* assays to investigate how functional and heterogenous CS granules form. Our findings support a model in which CS assembly begins with PCM1 scaffold formation, followed by regulated recruitment of clients. We identify mechanisms that control assembly, show that CS granules are spatially organized into subdomains and establish links between CS granule properties and cellular signaling. Collectively, our study introduces a conceptual framework and tools for studying CS homeostasis across diverse biological and disease contexts.

To dissect how PCM1 scaffolds CS assembly, we performed cellular reconstitution experiments with PCM1 deletion mutants. These showed that PCM1-M, which contains the second self-association domain, is sufficient for granule formation, whereas the C-terminal region fine-tunes CS properties rather than directly driving assembly. Together, these findings highlight PCM1 as a modular protein, with its C-terminal region regulating PCM1-N’s assembly activity, possibly through interactions with other proteins or post-translational modifications. This modularity may enable PCM1 to act as a molecular switch that initiates granule formation in response to cellular cues. Complementary evidence from in vitro reconstitution and induced dimerization assays further demonstrated that PCM1 multimerization is sufficient to drive scaffold formation and that its regulation is critical for CS organization and ciliary signaling. Thus, PCM1 multimerization emerges as the core principle of CS scaffold assembly, a mode of organization reminiscent of other multimerization-driven systems such as the scaffolding protein G3BP1 in stress granules ^73–75^.

To define the stages of the CS assembly pathway, we combined cellular biogenesis assays with 3D image analysis. Although this quantification cannot precisely resolve the absolute size of full-length PCM1 granules because they are near the diffraction limit, it allows robust relative comparisons across conditions. Using this approach, we found that assembly initiated at or near the centrosome, where PCM1 signal was first detected, followed by stochastic granule formation throughout the cytoplasm and subsequent pericentrosomal clustering. This pattern suggests that centrosomes may provide a favorable site for scaffold nucleation, possibly through microtubule-based enrichment of PCM1 above a critical threshold, although this remains to be tested. Analysis of scaffold dynamics further indicated a progressive transition toward a less dynamic state as assembly progressed.

Following PCM1 scaffold formation, clients are incorporated in a regulated manner that may contribute to CS organization and function. Co-localization analyses revealed that different clients associated with PCM1 at distinct times, suggesting a hierarchical process. These differences could underlie distinct client roles, with some regulating the scaffold and others modulating CS functions. For example, CEP131 and OFD1, which showed higher colocalization with PCM1 and were detected at earlier timepoints, may represent clients that associate closely with the scaffold and contribute to its regulation. In contrast, other proteins appeared later, likely being recruited as part of functional complexes or co-regulated groups. The BBSome, a seven-subunit complex that assembles at the CS before its ciliary targeting, illustrates this principle, with BBS4 identified as a late client in our assays ^76, 77^. Recruitment of different combinations of late proteins could further diversify CS composition and functions across contexts. We showed that client colocalization with PCM1 varied during ciliogenesis, providing evidence for a functional link between recruitment dynamics and CS activity. Hierarchical recruitment may also explain the emergence of distinct CS subtypes, depending on the initiating proteins and the sequence of recruitment events. Together, these findings support a model in which hierarchical client recruitment provides a potential mechanism for generating CS compositional heterogeneity and functional adaptability. Future studies, particularly time-resolved proteomic profiling, will be required to test this model systematically and to define the order and properties of recruited clients, as has been done for stress granules ^78, 79^.

We discovered internal organization of CS granules, which may provide an additional layer of regulation for their functions. High-resolution imaging revealed that CS granules are internally organized into distinct subdomains with different composition and dynamics. Clients showed varied spatial patterns: some followed a core–shell or radial distribution, while others occupied more irregular regions. PCM1 itself appeared internally partitioned, as N-, M-, and C-terminal probes localized to different regions within granules, suggesting that PCM1 contributes to internal organization. These subdomains also differed in their dynamics, with the PCM1 scaffold being relatively non-dynamic, while clients displayed varying levels of dynamic association. Such spatial organization, reminiscent of PML bodies, paraspeckles, and stress granules ^64, 80–84^, could allow co-regulated or functionally related proteins to cluster together within the same subdomain. In parallel, more dynamic clients that exchange with other cellular compartments may be concentrated in dynamic subdomains. ^64, 80–84^. Future studies using multiplex imaging and proteomic profiling of CS subdomains across contexts will be needed to define how internal organization contributes to their function.

In addition to the insights from cellular assays, in vitro reconstitution provided a powerful and complementary approach to test PCM1 behavior under defined conditions. Earlier studies using PCM1 fragments offered preliminary support for its intrinsic assembly capacity, but required non-physiological concentrations and additives ^27, 85^. By contrast, our experiments showed that full-length PCM1 forms granules at near-physiological levels and without crowding agents, allowing a more direct view of its properties. First, we demonstrated that homotypic interactions of PCM1 alone are sufficient to assemble a scaffold with internal organization. Second, PCM1 granules selectively recruited MIB1 and tubulin, and associated with microtubules. These findings are consistent with CS transport along microtubules and their functions in storing and trafficking clients. Finally, this reconstitution system provides a versatile platform to directly test the contributions of specific molecules such as kinases, RNA, or cellular extracts in regulating CS assembly and properties.

Together, our cellular and biochemical findings define the molecular principles of CS assembly and highlight PCM1 as the core scaffold for satellite organization and function. PCM1 mutations have been associated with schizophrenia, and C-terminal truncations that fuse with kinases have been identified in leukemia ^23, 86–91^. Moreover, mutations in several clients have been linked to disorders such as microcephaly and ciliopathies ^7, 13, 19^. By providing tools and a model for assembling CS granules, our work offers a framework to investigate how these mutations disrupt CS biogenesis and homeostasis, and how such alterations could contribute to disease mechanisms in neurodevelopmental disorders, ciliopathies, and cancer.

## Materials and Methods

### Plasmids

Full-length cDNAs of *Homo sapiens* PCM1 (GenBank accession no: NP_006188.4) and *Homo sapiens* MIB1 (GenBank accession no: NM_020774.4) were used. Primers used for cloning are included in Table S2. For PCM1, full-length cDNA and deletion mutants (nucleotides GFP-PCM1: 1-6072 bp, GFP-PCM1-N: 1:3600 bp and GFP-PCM1-M: 2100-3600 bp, GFP-PCM1-NS: 1:2100 bp and GFP-PCM1-C: 3600-6072 bp) were amplified by PCR and cloned into pDONR221 plasmid using Gateway recombination. ORFs of GFP-PCM1, GFP-PCM1-N and GFP-PCM1-M were amplified by PCR and cloned into pDONR221. Subsequent Gateway recombination reactions using pInducer20 and pEF-FRT-LapC DEST (provided by Nachury laboratory, UCSF) were performed. To generate a FLAG expression plasmid, PCM1 was cloned into the pCDNA5.1-FRT/TO-Flag-miniTurboID vector (provided by Bettencourt-Dias laboratory, IGC Gulbenkian) using AscI and NotI restriction sites.

For protein purification, cDNAs encoding GFP, mNeonGreen, PCM1-M (2100-3600 bp), and PCM1-C (4993-6073 bp) were amplified by PCR and ifirst cloned into pDONR221. These inserts were then transferred into pDEST-His-MBP and pDEST-GST expression plasmids by Gateway recombination. Full-length PCM1 was cloned into pCDH-EF1-mNeonGreen-T2A-Puro lentiviral plasmid using BamHI and NotI restriction sites. Full length MIB1 was cloned into pCDH-EF1-mScarlet-T2A-Puro lentiviral plasmid using the same restriction sites. mScarlet-MIB1 was amplified from the pCDH-mScarlet-MIB1 plasmid and cloned into the pDEST-His-MBP vector via Gateway recombination. mNeonGreen-tagged PCM1 was amplified from pCDH-mNeonGreen-PCM1 plasmid and cloned into the pFastBac-His-MBP vector by Gateway recombination.

For the inducible dimerization system, FKBP(F36V) was amplified from the pCDNA3.1(+)-miniTurboID-6xGGSGlinker-FKBPF36V-2xHA plasmid (Addgene plasmid #200641) and inserted into the pCDH-EF1-mNeonGreen-PCM1-T2A-Puro lentiviral plasmid using NotI restriction site.

### Cell culture and transfections

Human telomerase-immortalized retinal pigment epithelium cells (hTERT-RPE1, ATCC) were cultured in Dulbecco’s Modified Eagle’s Medium/F12 50/50 medium supplemented with 10% fetal bovine serum (FBS). Human embryonic kidney (HEK293T, ATCC) and human cervical carcinoma cells (HeLa, ATCC) were cultured in DMEM medium supplemented with 10% FBS. All mammalian cell lines were maintained in a humidified incubator at 37°C with 5% CO2 and confirmed to be negative for mycoplasma using a PCR-based assay. RPE1::PCM1 KO^10^, RPE1:mNeonGreen-CCDC66 ^35^ and RPE1:mNeonGreen-CSPP1 ^65^ cell lines were previously described and validated. Hela::PCM1 KO and HEK293T::PCM1 KO cells were generated using gRNA plasmids and CRISPR/Cas9 editing as previously described ^10^.

High Five (Hi-5) insect cells were cultured in serum-free Insect-XPRESS medium, while Spodoptera frugiperda (Sf9) insect cells were grown in Spodopan medium supplemented with 10% FBS for growth conditions. For transfections and baculovirus/protein production, Sf9 cells were switched to serum-free medium. Insect cell lines were maintained at 27°C with 5% CO_2_ in a humidified incubator, either as adherent monolayers or in suspension using a rotator shelf. Sf9 cells were transfected with plasmids using FuGENE HD for baculovirus production.

### Lentivirus production and cell transduction

Lentivirus was produced as previously described, using pInducer20-GFP-PCM1, pInducer20-GFP-PCM1-N, pInducer20-GFP-PCM1-M and pCDH-mNG-PCM1 FL-FKBP(F36V) plasmids as transfer vectors^92^. For infection, 1 × 10^5^ RPE1::PCM1 KO cells were seeded into 6-well tissue culture plates one day prior to transduction and infected the next day with 1 ml of viral supernatant. 24 h post-infection, the medium was replaced with fresh complete medium. After 48 h, the cells were split and subjected to antibiotic selection. Cells transduced with pInducer20-based lentivirus were selected in the presence of 1 mg/ml neomycin (G-418), while cells transduced with pCDH-mNG-based lentivirus were selected with 2 μg/ml puromycin. Selection was continued for 5-7 days until all uninfected control cells died

Infection efficiency in the heterogeneous pools was assessed by immunofluorescence. Subsequently, cells were trypsinized and serially diluted into normal growth medium to obtain single colonies. Colonies appearing after 10–14 days were picked, expanded, and screened for low-level expression of fusion proteins by immunofluorescence and Western blotting.

### Drug treatments

All chemicals used in this study are listed in Table S2.

For 1,6-hexanediol treatment, cells were incubated in complete medium containing 5% 1,6-hexanediol for 5 min, washed twice with PBS, and fixed. To induce stress granules, cells were treated with 50 mM NaAsO₂ for 1 h; their sensitivity to 1,6-hexanediol was then tested by subsequent incubation with 5% 1,6-hexanediol for 5 min.

For nocodazole treatment, inducible RPE1::PCM1 KO cells expressing GFP-PCM1 were subjected to two protocols: (i) for the 2 h assay, cells were treated with 5 μg/ml nocodazole for 1 h in parallel with induction by 1 μM doxycycline for 2 h; (ii) for the 16 h assay, cells were simultaneously induced with 1 μM doxycycline and treated with 5 ng/ml nocodazole for 16 h. Following treatment, cells were washed with PBS and fixed.

For RNase treatment, cells were incubated in complete medium containing 100 μg/ml RNase I and RNase A for 1 h, then washed with PBS and fixed.

For ciliation assays, cells were serum-starved in medium containing 0.5% FBS for 24 or 48 h after reaching full confluency. For Hedgehog pathway activation, serum-starved cells were treated with 50 nM SAG for 24 h.

For inducible dimerization of PCM1, RPE1::PCM1 KO cells stably expressing mNG-PCM1-FKBP(F36V) were treated with 250 nM B/B Homodimerizer (AP20187) for at least 4 h. To induce dissociation of dimerized granules, cells were washed twice with PBS and subsequently treated with 1 μM wash-out ligand for a minimum of 1 h. Treatment duration varied according to the experiment: 24 h for cilium assembly, 48 h for Sonic hedgehog stimulation, and 18 h for live-cell imaging of mitosis.

### Immunofluorescence, antibodies and microscopy

Cells were grown on coverslips, washed twice with PBS, and fixed with ice-cold methanol at −20 °C for either 3 or 10 min. After two additional PBS washes, cells were blocked with 3% BSA in PBS containing 0.1% Triton X-100 and incubated with primary antibodies diluted in blocking solution for 1 h at room temperature. Cells were then washed three times with PBS, incubated with secondary antibodies and DAPI (1:2000) for 45 min at room temperature, washed again three times with PBS, and mounted in Mowiol mounting medium containing N-propyl gallate. A complete list of primary and secondary antibodies used for immunofluorescence experiments is provided in Table S2.

Confocal microscopy was performed on a Leica TCS SP8 or Stellaris 8 laser-scanning inverted confocal microscope equipped for four-color imaging with 405, 488, 561, and 633 nm laser lines; four detection channels (one photomultiplier tube and three hybrid detectors); HC PL APO CS2 63×/1.4 NA and 40×/1.3 NA oil objectives; and an incubation chamber. The pinhole was set to 1 µm. Images were acquired as z-stacks of 20–50 optical sections with 0.2–0.5 µm spacing, using line averaging of 2–3 and a 400 Hz unidirectional xyz scan mode. Images were collected at 1,024 × 1,024 pixel resolution with an additional 2–4× optical zoom as needed. Acquisition was controlled with Leica Application Suite X (LAS X) software, and post-processing was performed using either Huygens Professional (Scientific Volume Imaging) or LAS X Lightning.

3D-SIM imaging was carried out on an Elyra 7 system with Lattice SIM2 (Carl Zeiss Microscopy) using a Plan-Apochromat 63×/1.4 Oil DIC M27 objective, 405/488/561/633 nm laser illumination, and standard excitation/emission filter sets. Images were acquired with a sCMOS (version 4.2 CL HS) camera at 110 nm z-spacing over a 5–10 µm thickness. During acquisition, laser powers at the focal plane were 2– 12%, exposure times were 50–250 ms, and camera gain values were 5–50. Raw images were reconstructed using the SIM module of ZEN Black software. Channel alignment was performed using calibration files generated from super-resolution Tetraspec beads (Carl Zeiss Microscopy).

Time-lapse imaging was performed with doxycycline-inducible RPE1::PCM1 KO cell lines expressing GFP-PCM1, GFP-PCM1-N, or GFP-PCM1-M, seeded on 35-mm glass-bottom dishes. Following induction with 1 µg/ml doxycycline, live imaging was carried out on a Stellaris 8 confocal microscope equipped with an HC PL APO CS2 63×/1.4 NA oil objective, maintained at 37 °C with 5% CO₂. The same acquisition parameters were used for replicate experiments. For time-lapse imaging of CS dimerization and dissociation, cells were cultured in Ibidi µ-Slide 8-well chambers and imaged at 37 °C and 5% CO₂. Images were acquired every 2 min for the first hour, followed by every 20 min up to 4 h, using a 0.5 µm z-step size and a 512 × 512 pixel format. Dimerizer and washout ligands were applied between the first two frames.

### Ultrastructural Expansion Microscopy

Ultrastructure Expansion Microscopy (U-ExM) was performed as previously described^93^. RPE1 cells were cultured on 12 mm coverslips in 24-well plates and fixed in 1.4% formaldehyde/2% acrylamide (2× FA/AA) in 1× PBS for 5 h at 37 °C. Cells were then embedded in a gel prepared from Monomer Solution (38% sodium acrylamide [w/w], 12.5 µl acrylamide, 2.5 µl BIS, and 5 µl 10× PBS), supplemented with TEMED and 0.5% APS, and polymerized for 1 h at 37 °C.

Denaturation was carried out at 95 °C for 90 min, followed by staining with primary antibodies for 3 h at 37 °C. Gels were washed three times for 10 min each in PBS-T at room temperature (RT), incubated with secondary antibodies for 2.5 h at 37 °C, and washed again three times for 10 min in PBS-T at RT.

Before imaging, gels were expanded three times in 100 ml dH₂O. The expansion factor was determined by measuring the gel diameter with a ruler and dividing it by the original 12 mm coverslip diameter. Expanded gels were cut into smaller pieces and mounted on 24 mm poly-D-lysine–coated coverslips. Imaging was performed on a Leica Stellaris confocal microscope using 0.30 µm z-intervals, and images were deconvolved with Lightning mode.

### Image Analysis

#### 1. Quantification of granule properties and threshold concentrations

Granule properties, including number, size, and intensity, were quantified from GFP-PCM1 or endogenous PCM1 signals using the Surface Creation function in Imaris software (Bitplane, Oxford Instruments). Surfaces were detected by intensity thresholding, and diameters were estimated using the software’s built-in algorithm. Manual corrections were applied as needed to ensure accurate segmentation. Centrosomes were analyzed with the Spot Detection module in Imaris, based on γ-tubulin or centrin 2 signals. Spot size was set to ∼1 µm, and automatic detection was performed with an intensity threshold. False positives were removed by manual curation. For each cell, parameters including volume (µm³), average volume (µm³), intensity sum, average intensity, distance to spots, sphericity, and surface count were exported as Excel files and analyzed in GraphPad Prism.

Threshold concentrations were determined from relative protein concentrations, calculated using total fluorescence intensities and volumes. PCM1 granules were segmented from the GFP signal without applying the smoothing option, and the *Split Touching Objects* function was used to separate adjoining granules. Different thresholds were applied to distinguish granular from cytoplasmic PCM1. Both were rendered in 3D, and total intensity and volume values were exported. Intensity values were normalized to the maximum fluorescence intensity for each biological replicate. Granule concentration was calculated by dividing total intensity by total volume (µm³) per cell. Cytoplasmic PCM1 concentration was obtained by subtracting the granule intensity and volume from the whole-cell values, and dividing the remaining ratio. The threshold concentration of PCM1 granules was defined as the average concentration across all granules in each cell line.

Colocalization was quantified using Pearson’s correlation coefficient in the Imaris *Coloc* module. Endogenous PCM1 or GFP-PCM1 was assigned as channel 1, and client proteins as channel 2. Thresholding parameters were chosen to specifically identify PCM1 granules, and a new channel was generated using the *Build Coloc Channel* function. Results were exported via the Image Properties tab and analyzed in GraphPad Prism.

For *in vitro* experiments, granule quantification was performed using Aivia software (Leica Microsystems). Prior to analysis, raw 3D image stacks were background-subtracted to improve segmentation by the machine learning tool. Granules were detected using the Object Detection module with optimized intensity thresholding and size filtering to exclude non-specific signals. Segmentation parameters were fine-tuned to eliminate background objects and accurately detect individual granules. Granule volume values were exported and analyzed in GraphPad Prism, with radii calculated assuming spherical shape.

#### 2. Line profile calculations

3D images were opened in ImageJ in their native format, and z-stacks were projected using the *3D Project* function (Image > Stacks > 3D Project). A line of varying length was drawn across granules, and intensity values along the line were measured with *Analyze > Plot Profile*. Exported values were plotted in GraphPad Prism.

#### 3. Analysis of centrosomal and ciliary protein levels

Centrosomal and ciliary protein levels were quantified in ImageJ. Regions of interest (ROIs) were drawn around centrosomes (marked by γ-tubulin or centrin 2) or primary cilia (marked by Arl13b, acetylated tubulin, or glutamylated tubulin). Corresponding signals were measured, and cytoplasmic background was subtracted for each cell. Values were normalized to the mean intensity of each biological replicate for statistical analysis.

### Fluorescence recovery after photobleaching analysis

RPE1::PCM1 KO stable cell lines expressing GFP-PCM1-N or GFP-PCM1-M were cultured in glass-bottom dishes (Lab-Tek II Chambered Coverglass) and induced with 1 µM doxycycline for 20 min, 1 h, or 16 h. Photobleaching was performed on a Leica SP8 confocal laser scanning microscope equipped with a Plan-Apo 63×/1.4 NA oil objective and a scan zoom of 2.5. Cells were maintained at 37 °C with 5% CO₂ throughout imaging. z-stacks spanning 2.7 µm with a 0.3 µm step size were acquired before and after bleaching. A single pre-bleach image stack was captured, followed by bleaching of defined ROIs using a 488-nm Argon laser at 100% power for 10 iterations. Post-bleach image stacks were acquired every 8 s for 5 min. Relative fluorescence intensities were calculated from maximum intensity projections using Fiji, and plotted as a function of time. Quantification of recovery curves, half-time (t½), and mobile fractions was performed using previously described equations^94^.

### Cell lysis and immunoblotting

Cells were lysed in RIPA buffer containing 50 mM Tris-HCl, 150 mM NaCl, 1.0% (v/v) Triton X-100, 0.5% (w/v) sodium deoxycholate, 1.0 mM EDTA, 0.1% (w/v) SDS, and protease inhibitors. Lysis was carried out for 30 min at 4°C, followed by centrifugation at 13,000 g for 15 min. Protein concentration in the supernatant was measured using the Bradford assay (Bio-Rad Laboratories). For immunoblotting, equal amounts of proteins were loaded and separated by SDS-PAGE, transferred onto nitrocellulose membranes, and blocked with 5% milk in TBS-T for 1 h at room temperature. Membranes were incubated overnight at 4°C with primary antibodies diluted in 5% BSA in TBS-T, followed by three 5-minute washes in TBS-T. They were then incubated with secondary antibodies for 1 h at room temperature, washed again, and visualized using the LI-COR Odyssey Infrared Imaging System at 680 and 800 nm (LI-COR Biosciences). Full list of primary antibodies used for western blot experiments in this study are listed Table S2. Secondary antibodies used for Western blotting experiments were IRDye 680- and IRDye 800-coupled and were used at 1:15,000 (LI-COR Biosciences).

### Solubility assay

HEK293T::PCM1 KO cells were co-transfected with GFP-PCM1, GFP-PCM1-N, GFP-PCM1-C, GFP-PCM1-M, or GFP-PCM1-NS. After 48 h, cells were lysed in buffer containing 50 mM HEPES (pH 8), 100 mM KCl, 2 mM EDTA, 10% glycerol, 0.1% Nonidet™ P40, and a protease inhibitor cocktail. Lysates were incubated on ice for 30 min and centrifuged at 14,000 rpm for 15 min at 4 °C. Protein concentrations in the supernatants were measured using the Bradford assay. Both the soluble supernatant and the insoluble pellet fractions were resuspended in SDS sample buffer and analyzed by immunoblotting.

### Pulldown experiments

For FLAG pulldown experiments, HEK293T::PCM1 KO cells were co-transfected with Flag-PCM1 together with GFP, GFP-PCM1-N, GFP-PCM1-C, GFP-PCM1-M, or GFP-PCM1-NS. After 48 h, cells were washed and lysed in FLAG pulldown buffer (50 mM HEPES, pH 8; 100 mM KCl; 2 mM EDTA; 10% glycerol; 0.1% NP-40; freshly supplemented with 1 mM DTT and protease inhibitors: 10 µg/ml leupeptin, pepstatin, and chymostatin, plus 1 mM PMSF) for 30 min on ice. Lysates were centrifuged at 13,000 rpm for 10 min at 4 °C, and supernatants were collected. A 100 µl aliquot of each sample was reserved as input, and the remaining supernatant was incubated overnight at 4 °C with Anti-FLAG M2 agarose beads for immunoprecipitation.

### Protein expression and purification

For the expression of His-MBP-mNG-PCM1, 500 ml of Hi5 cells at a density of 1 × 10⁶ cells/ml were infected with P1/P2 baculovirus produced in Sf9 cells at an MOI of 1 and harvested 48 h post-infection. For the expression of His-MBP-mScarlet-MIB1, BL21 Rosetta cells were induced at an OD_600_ of 0.5 with 0.5 mM IPTG, incubated at 18 °C for 18 h, and then harvested.

For protein purification, cells were lysed in ice-cold lysis buffer (50 mM NaH₂PO₄, 500 mM NaCl, 20 mM imidazole, 0.5% Triton, 10% glycerol, 0.5 mM arginine, pH 7.0) supplemented with 1 mM PMSF, 5 mM β-mercaptoethanol, and a protease inhibitor cocktail. Lysates were sonicated and clarified by centrifugation at 14,000 rpm for 45 min at 4 °C. Proteins were bound to Ni-NTA beads, eluted with 300 mM imidazole, and dialyzed overnight at 4 °C into storage buffer.

MIB1 was dialyzed into buffer containing 20 mM HEPES (pH 7.0), 500 mM NaCl, and 1 mM DTT. PCM1 was dialyzed into the same buffer using a PD-10 desalting column. Proteins were then concentrated with Amicon Ultra 30K centrifugal filter devices. Finally, 10 μl aliquots were snap-frozen in liquid nitrogen and stored at −80 °C.

### *In vitro* PCM1 granule reconstitution

Purified His-MBP-mNG-PCM1 was diluted to 0.5 μM, 1 μM or 2 μM in low-salt assay buffer (10 mM HEPES, 0.1 mM EDTA, 2 mM DTT, 150 mM NaCl). After a 5 min incubation, 25 μL of each mixture was loaded into an imaging chamber constructed by mounting HCl-treated glass coverslips onto slides with strips of double-sided tape, as previously described ^95^. Imaging was performed using a Leica SP8 confocal microscope with an HC PL APO CS2 63× 1.4 NA oil objective. Additional reactions containing 0.15 μM His-MBP-mScarlet-MIB1 were analyzed under the same buffer, setup and imaging conditions, where His-MBP-mScarlet-MIB1 was added to the reaction mix..

For examining microtubule association and the regulation of CS by microtubules, 1 mM Taxol-stabilized microtubules were analyzed in BRB80 buffer (80 mM PIPES pH 6.8, 1 mM EGTA, 1 mM MgCl2) using the same setup. To generate taxol-stabilized microtubules, purified bovine tubulin was first precleared at 90,000 rpm for 5 min at 4 °C using a Beckman Coulter Optima MAX-XP ultracentrifuge with a TLA110 rotor. The cleared tubulin was diluted to 2 mg/mL and polymerized at 37 °C for 30 min in the presence of 20 μM Taxol added stepwise at 0.02, 0.2, and 2 μM. After polymerization of the microtubules, purified His-MBP-mNG-PCM1 granule reconsitiution assays were performed in BRB80 buffer to maintain the microtubule integrity for with and without microtubule conditions by mixing microtubules and purified His-MBP-mNG-PCM1 in BRB80 buffer. After a 5 min incubation, 25 μL of each mixture was loaded into an imaging chamber constructed by mounting HCl-treated glass coverslips onto slides with strips of double-sided tape.

For TIRF analysis, HCl-treated coverslips mounted onto slides with double-sided tape. Rhodamine-labeled microtubules (1 μM) were polymerized from bovine brain tubulin containing 10% Alexa Fluor™ 568–labeled tubulin mixed with 90% unlabeled tubulin. Microtubules were immobilized on silanized coverslips coated with anti–β-tubulin antibody, prepared by trichloroethylene/dichloromethylsilane treatment. Excess antibody was blocked with 0.2 mg/mL κ-casein and Pluronic-127. After attachment of the microtubules to the slides, microtubules were sequentially washed with 2 mg/mL κ-casein and then 0.05 mg/mL κ-casein. A solution containing 140 nM His-MBP–mNG-PCM1, κ-casein, and an oxygen scavenger system (glucose oxidase, catalase, β-mercaptoethanol, and glucose in BRB80) was introduced and incubated for 3 min to allow PCM1 binding to microtubules. Samples were sealed with nail polish and imaged at 30 °C on a Zeiss ELYRA 7 microscope using a 100× TIRF objective.

### Statistical analysis

Statistical significance and p-values were assessed using Prism 7 software (GraphPad Software, La Jolla, CA). For comparisons between two groups with a normal distribution, a two-tailed Student’s t-test was used, while one-way analysis of variance (ANOVA) was applied for comparisons involving more than two groups. The sample size (n) represents the number of experimental replicates for each sample and condition. Error bars in the graphs indicate the standard deviation (SD), and statistical significance levels were denoted as follows: ns: P>0.05, *:P<0.05, **:P<0.01, ***:P<0.001 and ****:P<0.0001.

## Acknowledgements

We thank the members of CytoLab for their insightful feedback on this work. We are grateful to Dila Gulensoy for cloning plasmids to express GFP fusions of PCM1-NS, PCM1-N, and PCM1-C, and to Dr. Jovana Deretic for assistance with TIRF experiments. We also acknowledge the use of services and facilities at the Koç University Animal Facility and the Istanbul University Animal Facility. Support for the use of Imaris software was provided by the Emory University Integrated Cellular Imaging Core Facility (RRID:SCR_023534).

This project received funding from the European Union’s Horizon Europe research and innovation program under the European Research Council Starting Grant “SatelliteHomeostasis” awarded to ENF. Additional support was provided by the EMBO Installation Grant (3622), the EMBO Young Investigator Award, and a grant from the Istanbul Development Agency, all awarded to ENF.

## Competing interest

The authors declare no competing interests.

## Notes

### Competing Interest Statement

The authors have declared no competing interest.

### Summary of Updates

This revised version includes substantial editorial improvements to enhance readability and clarity. We refined the text throughout, removed some data for conciseness and revised several figures to improve visual clarity and consistency. In addition, we reorganized certain sections to better highlight our main findings.

