## Supplementary Figures for "Centriolar satellites assemble via a hierarchical pathway driven by PCM1 multimerization"

### Figure S1

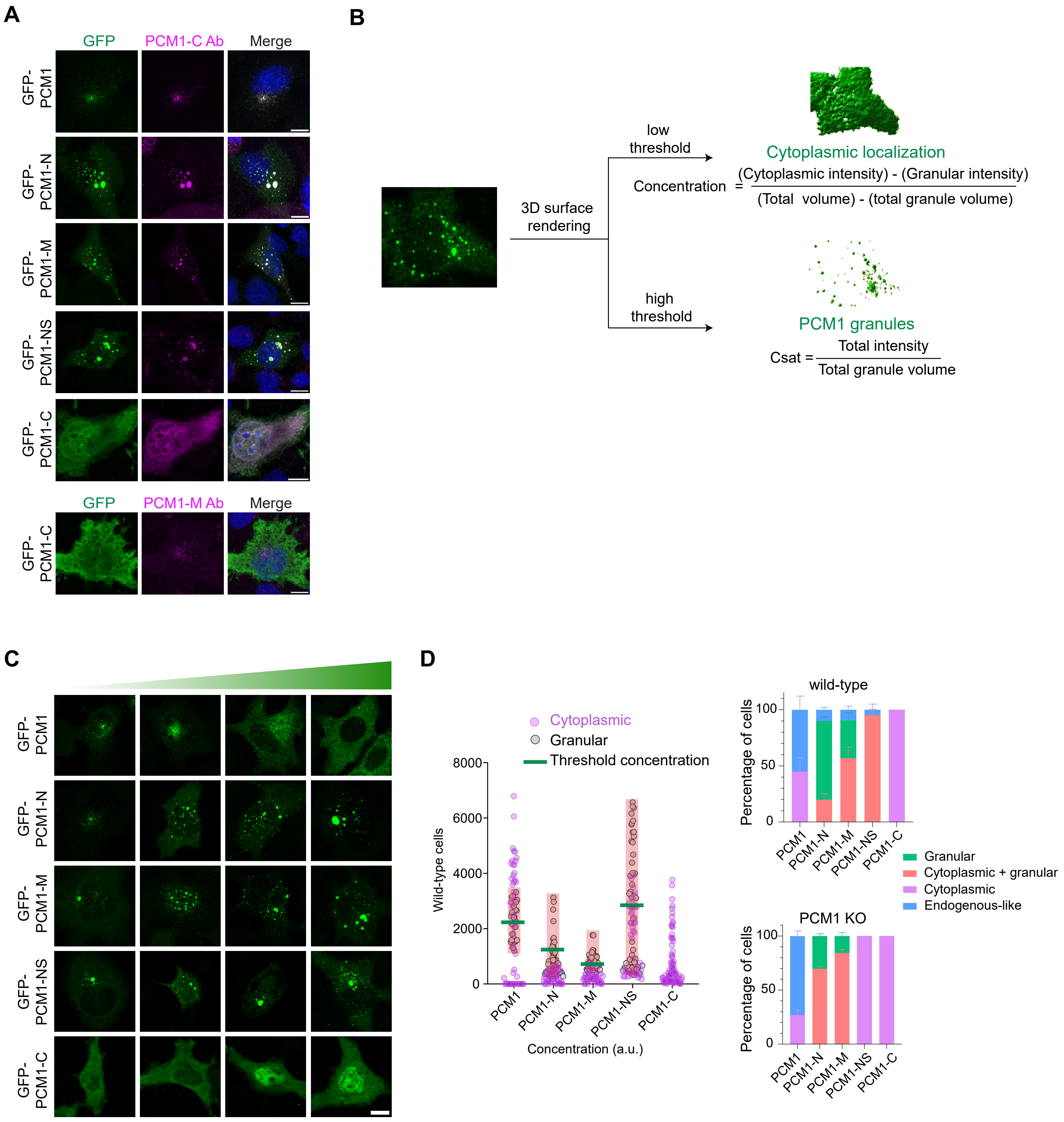

A
GFP
PCM1-C Ab
Merge
GFP-PCM1
GFP-PCM1-N
GFP-PCM1-M
GFP-PCM1-NS
GFP-PCM1-C
GFP
PCM1-M Ab
Merge
GFP-PCM1-C
B
3D surface rendering
low threshold
Cytoplasmic localization
Concentration = (Cytoplasmic intensity) - (Granular intensity) / (Total volume) - (total granule volume)
high threshold
PCM1 granules
Csat = Total intensity / Total granule volume
C
GFP-PCM1
GFP-PCM1-N
GFP-PCM1-M
GFP-PCM1-NS
GFP-PCM1-C
D
Cytoplasmic
Granular
Threshold concentration
Wild-type cells
8000
6000
4000
2000
0
PCM1
PCM1-N
PCM1-M
PCM1-NS
PCM1-C
Concentration (a.u.)
wild-type
Percentage of cells
100
50
0
PCM1
PCM1-N
PCM1-M
PCM1-NS
PCM1-C
Granular
Cytoplasmic + granular
Cytoplasmic
Endogenous-like
PCM1 KO
Percentage of cells
100
50
0
PCM1
PCM1-N
PCM1-M
PCM1-NS
PCM1-C

**Supplementary Figure S1**

**Characterization of distinct PCM1 domains**

**(A)** Recruitment of endogenous PCM1 to fragment-induced granules. Representative confocal images of HeLa-WT cells transfected with GFP-PCM1 and its truncations. Cells are stained with anti-GFP (green), PCM1-C-Ab (magenta) or PCM1-M-Ab (magenta) and DAPI (blue). GFP-PCM1-NS, GFP-PCM1-N and GFP-PCM1-M recruit endogenous PCM1 into granules. Similar results were obtained from three biological replicates. Scale bars: 10 µm.

**(B)** Workflow of determining the relative protein concentration and critical saturation concentration. The schematic illustrates the process of transforming z-stack confocal images into three-dimensional voxels using Imaris software. This transformation was performed for two different volumetric representations: one combining both cytoplasmic and condensate volumes, and the other representing only the condensate volumes. Following rendering, the sum of intensities and the volume of the 3D voxels were quantified. These measurements were subsequently used to determine the relative protein concentrations within the light and dense phases.

**(C)** Localization of PCM1 and its truncation mutants at increasing concentrations in HeLa-WT cells. These cells are transfected with GFP fusions of PCM1, PCM1-N, PCM1-M, PCM1-NS and PCM1-C for 48 hours, fixed and stained with anti-GFP. The images indicate the cells with increasing expression levels of PCM1 and its truncations, indicated by a green triangle at the top.

**(D)** Quantification of the threshold concentrations at which granules formed for PCM1 and its truncations from (C). For each transfected cell, intensity of GFP signal and total volume and intensity of granules and cytoplasm were measured using Imaris Software. Threshold concentration denotes the mean value of granular concentration values. Results are from three biological replicates, min 48 cells in total. a.u.: arbitrary unit. Quantification of the percentage of HeLa-WT and HeLa::PCM1 KO cells exhibiting distinct PCM1 localization patterns: granular, cytoplasmic, granular+cytoplasmic, and endogenous-like. Granular cells display no cytoplasmic localization and exhibit higher fluorescence intensity than the background signal. Endogenous-like refers to granules similar to endogenous CS granules, whereas granular describes granules that are larger and more heterogeneous in number. Data represent the mean percentage of cells from three biological replicates, totaling 100 cells. Scale bar: 10 µm.

Figure S2

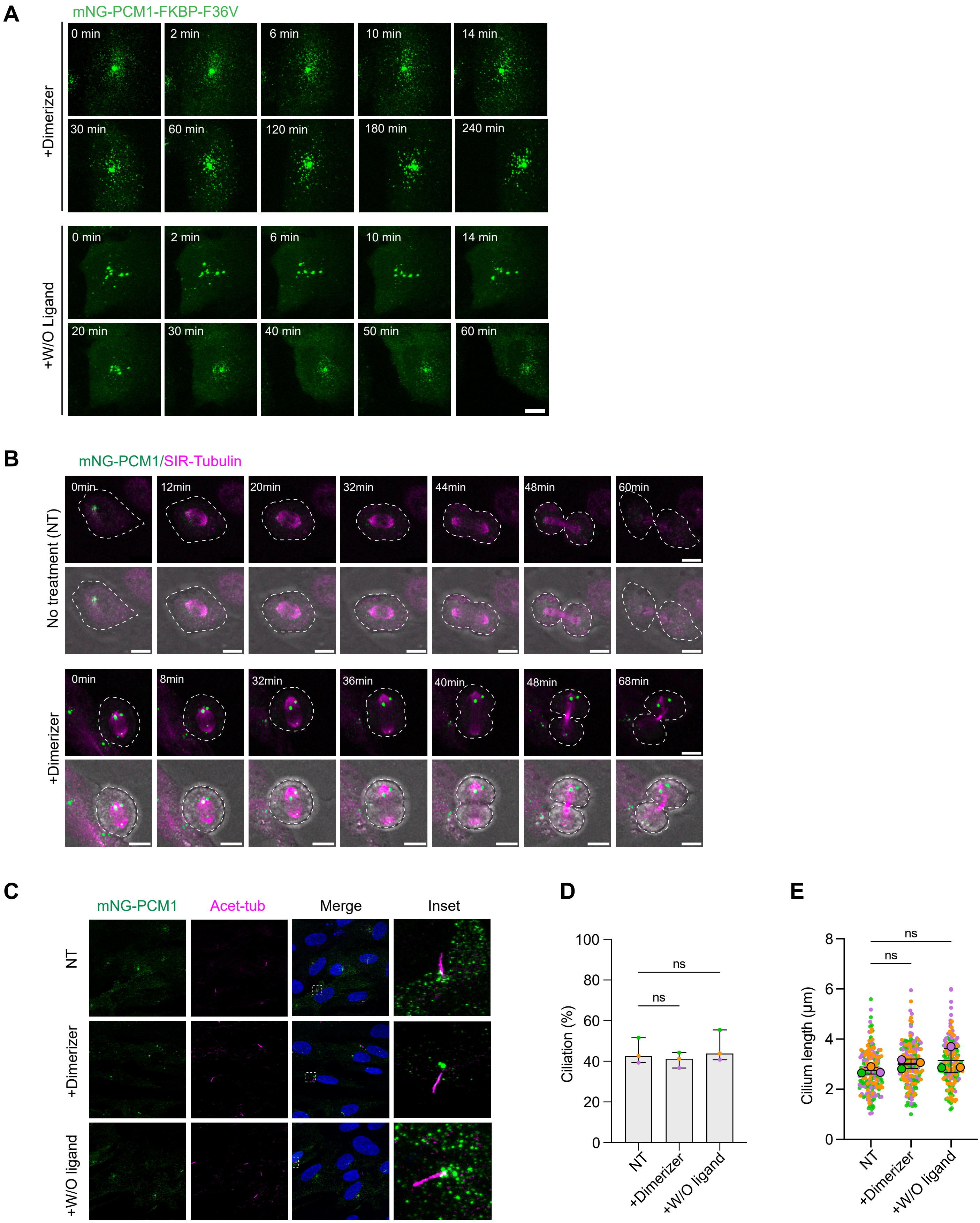

A
mNG-PCM1-FKBP-F36V
+Dimerizer
0 min
2 min
6 min
10 min
14 min
30 min
60 min
120 min
180 min
240 min
+W/O Ligand
0 min
2 min
6 min
10 min
14 min
20 min
30 min
40 min
50 min
60 min
B
mNG-PCM1/SIR-Tubulin
No treatment (NT)
0min
12min
20min
32min
44min
48min
60min
+Dimerizer
0min
8min
32min
36min
40min
48min
68min
C
mNG-PCM1
Acet-tub
Merge
Inset
NT
+Dimerizer
+W/O ligand
D
Ciliation (%)
100
80
60
40
20
0
ns
ns
NT
+Dimerizer
+W/O ligand
E
Cilium length (µm)
8
6
4
2
0
ns
ns
NT
+Dimerizer
+W/O ligand

**Supplementary Figure S2**

**Characterization of CS spatiotemporal dynamics and function in the chemically inducible dimerization system.**

**(A)** Still images depicting the behavior of CS granules upon treatment with dimerizer or washout ligand in RPE1::PCM1-KO cells stably expressing mNG-PCM1-FKBP(F36V). Live imaging was conducted at one-minute intervals for the first hour, followed by 20-minute intervals thereafter. Scale bar: 30 µm.

**(B)** CS granules after inducible dimerization persist through mitosis and segregate randomly into daughter cells. Still images from the time-lapse movie shows CS granule (igreen) behavior during mitosis in both control and dimerizer-treated cells. Both conditions were also treated with SiR-tubulin to visualize spindle microtubules (magenta). Live imaging was conducted at 5 minute intervals for 18 hours. Scale bar: 10 µm.

**(C)** Effects of CS dimerization on ciliogenesis and cilium length. Representative images depict ciliated RPE1::PCM1 KO cells stably expressing mNG-PCM1-FKBP(F36V) after treatment with dimerizer and subsequent wash-out ligand in serum-depleted medium for 24 hours. Cells were fixed and stained for mNeonGreen and acetylated tubulin. Scale bars: 20 µm (main panels), 5 µm (insets).

**(D, E)** Quantification of (D) ciliation percentage and (E) cilium length in control, dimerizer-treated, and wash-out ligand-treated cells 24 hours after serum starvation. The data represent three biological replicates, green, magenta and orange colored dots representing different replicates, with n = 35 per replicate.

#### Figure S3

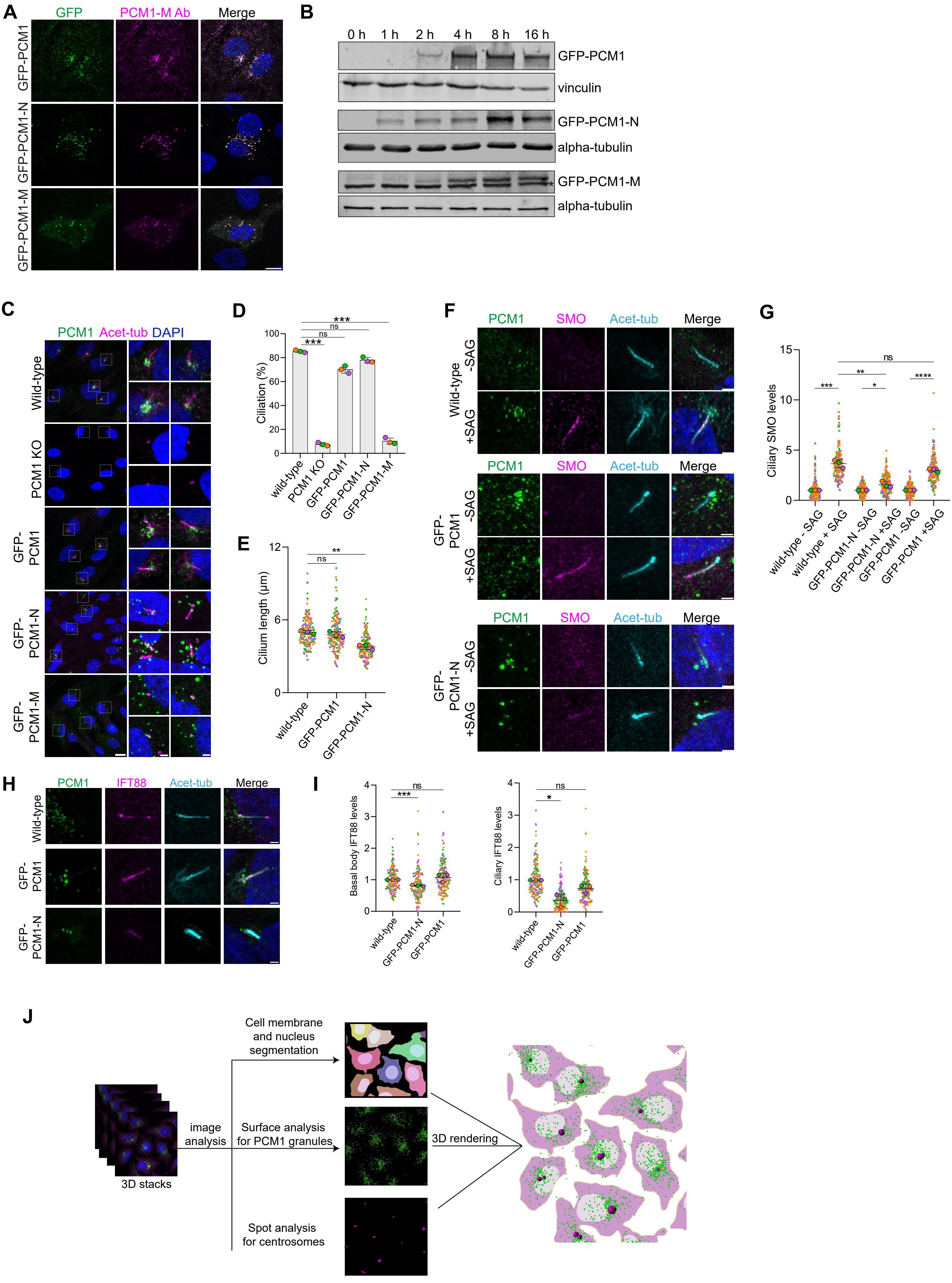

A
GFP
PCM1-M Ab
Merge
GFP-PCM1
GFP-PCM1-N
GFP-PCM1-M
B
0 h
1 h
2 h
4 h
8 h
16 h
GFP-PCM1
vinculin
GFP-PCM1-N
alpha-tubulin
GFP-PCM1-M
alpha-tubulin
C
PCM1 Acet-tub DAPI
Wild-type
PCM1 KO
GFP-PCM1
GFP-PCM1-N
GFP-PCM1-M
D
Ciliation (%)
wild-type
PCM1 KO
GFP-PCM1
GFP-PCM1-N
GFP-PCM1-M
E
Cilium length (µm)
wild-type
GFP-PCM1
GFP-PCM1-N
F
PCM1
SMO
Acet-tub
Merge
Wild-type
-SAG
+SAG
GFP-PCM1
GFP-PCM1-N
G
Ciliary SMO levels
wild-type - SAG
wild-type + SAG
GFP-PCM1-N -SAG
GFP-PCM1-N +SAG
GFP-PCM1 -SAG
GFP-PCM1 +SAG
H
PCM1
IFT88
Acet-tub
Merge
Wild-type
GFP-PCM1
GFP-PCM1-N
I
Basal body IFT88 levels
Ciliary IFT88 levels
wild-type
GFP-PCM1-N
GFP-PCM1
J
3D stacks
image analysis
Cell membrane and nucleus segmentation
Surface analysis for PCM1 granules
Spot analysis for centrosomes
3D rendering

**Supplementary Figure S3**

**Generation and validation of inducible cells for *de novo* CS biogenesis assay and their functional characterization in ciliogenesis and signaling.**

**(A)** Validation of inducible cell lines by immunofluorescence. Representative confocal images of RPE1::PCM1 KO cells inducibly expressing GFP-PCM1, GFP-PCM1-N and GFP-PCM1-M fixed and stained after 16 hours of induction. Cells were stained with anti-GFP-Ab (green), anti-PCM1-M-Ab (magenta) and DAPI (blue). Scale bar: 5 μm.

**(B)** Validation of inducible cell lines by immunoblotting. Immunoblot analysis of RPE1::PCM1 KO inducible cell lines expressing GFP-PCM1, GFP-PCM1-N and GFP-PCM1-M. Cell lines were induced with doxycycline for 1, 2, 4, 8 and 16 hour and lysates were prepared along with non-induced control (0 h). Proteins were detected using anti-GFP antibody and either vinculin or alpha-tubulin as loading controls. Similar results were obtained from three biological replicates. *non-specific band.

**(C)** Experiments to assess phenotypic rescue in ciliation efficiency and cilium length. RPE1::PCM1 KO cells inducibly expressing GFP-PCM1, GFP-PCM1-N and GFP-PCM1-M were induced by doxycycline for 48 hours, then serum starved for 48 hours. RPE1-WT and RPE1::PCM1 KO cells were serum starved for 48 hours. Cells were fixed and stained with anti-GFP (green), anti-acetylated tubulin (magenta), and DAPI for nucleus(blue). The percentage of ciliated cells was determined from acetylated tubulin signals. Scale bars: 10 μm (main panels), 2 μm (insets).

**(D)** Quantification of ciliation percentage from (G). Only GFP-positive cells were chosen for the quantification. Data represents the mean from n>40 cells per replicate and three biological replicates. ***: P<0.001; ns: not significant.

**(E)** Quantification of cilium length from (G). Only GFP-positive cells were chosen for the quantification. Data represents the mean from n>40 cells per replicate and three biological replicates, green, magenta and orange colored dots representing different replicates. ***: P<0.001; ns: not significant.

**(F)** Experiments to assess phenotypic rescue in Hedgehog pathway activation. RPE1::PCM1 KO cells, which inducibly express GFP-PCM1 and GFP-PCM1-N, were treated with doxycycline for 48 hours, then serum-starved for 24 hours, and subsequently treated with 500 nM SAG in serum starvation medium for 24 hours. In parallel, RPE1-WT cells were serum-starved for 24 hours and subsequently treated with 500 nM SAG for 24 hours. Basal activation levels were determined by processing control cells that were not treated with SAG (SAG-). All cells were fixed and stained for GFP (green), Smo (magenta), acetylated tubulin (cyan), and DAPI (blue). Scale bars: 2 μm.

**(G)** Quantification of Smo recruitment to the cilium as shown in (J). Smo intensity at the cilium was calculated using ImageJ. Only GFP-positive cells were selected for quantification. The data represent the mean from three biological replicates, with n>55 cells per replicate, green, magenta and orange colored dots representing different replicates. *:P<0.05, **:P<0.01, ***:P<0.001, ****:P<0.0001; ns: not significant.

**(H)** Experiments to assess phenotypic rescue in recruitment of IFT88 to the cilium and basal body. RPE1::PCM1 KO cells inducibly expressing GFP-PCM1 and GFP-PCM1-N were treated with doxycycline for 48 hours, then serum-starved for 48 hours. In parallel, RPE1-WT cells were serum-starved for 48 hours. All cells were fixed and stained for GFP (green), IFT88 (magenta), acetylated tubulin (cyan), and DAPI (blue). Scale bar: 2 µm.

**(I)** Quantification of ciliary and centrosomal IFT88 levels from (L). The results shown represent the mean of three independent experiments. The basal body signal was measured from a 3 $\mu m^2$ area centered on the centrosome, as identified by the acetylated tubulin signal. Data are presented as the mean from three biological replicates, with n>55 cells per replicate, green, magenta and orange colored dots representing different replicates.

**(J)** Workflow for quantifying CS granule properties. PCM1 granule characteristics were analyzed in both fixed and live-cell images using Imaris software. The analysis was conducted in 3D, with quantifications performed post 3D rendering. Cells were segmented individually per the 'one cell per nucleus' rule using the cell creation tool. The surface creation tool was used to render PCM1 granules, incorporating background subtraction and manual thresholding for granule detection. Centrosomes were identified using centrin-2 or gamma tubulin channels using the spot creation tool, set to a 1 µm XY diameter. All three segmented components were combined to form a 3D rendered image, from which granule properties (volume, fluorescence intensity, average distance from the centrosome) were quantified.

### Figure S4

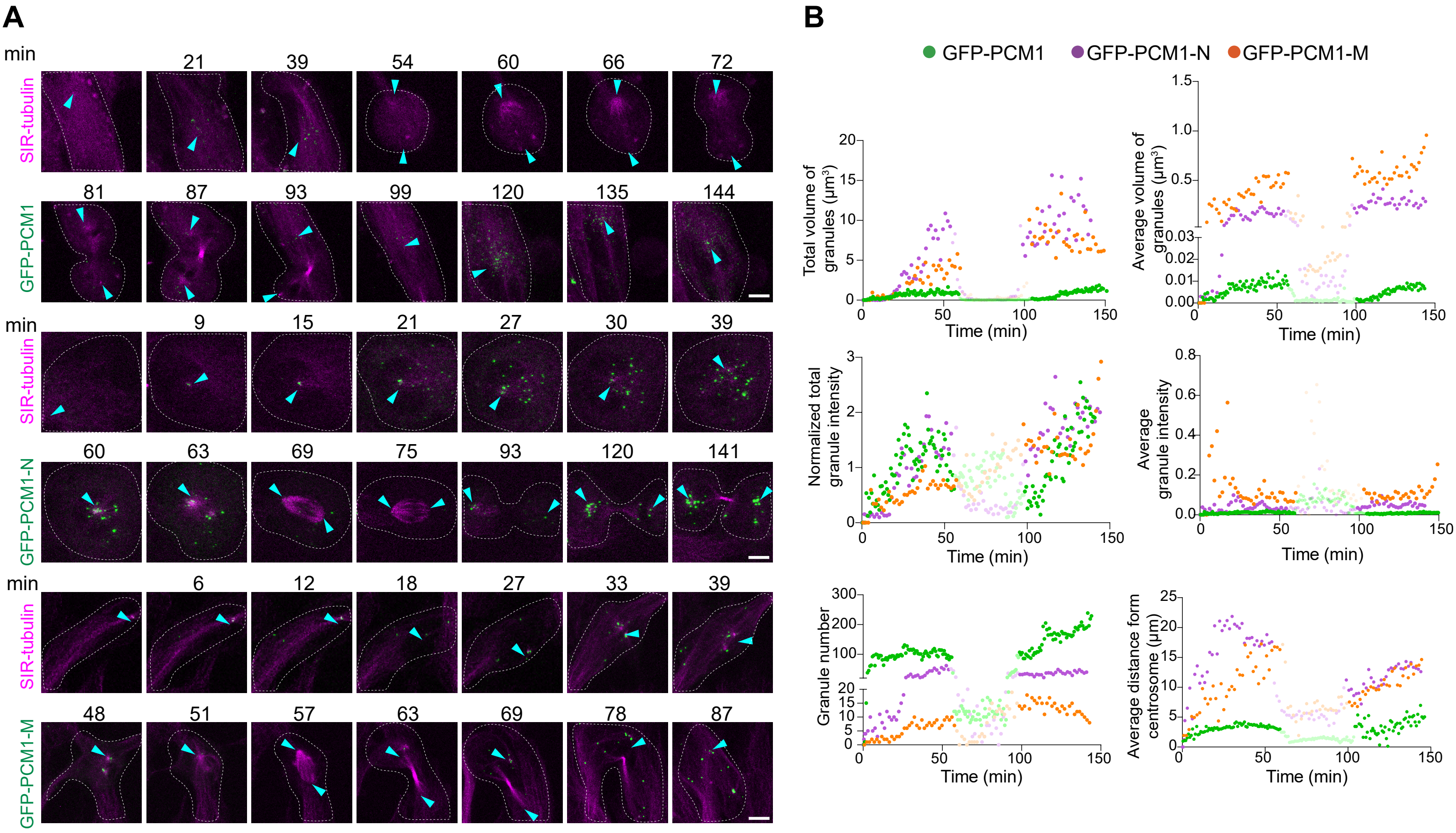
A
B
min
SIR-tubulin
GFP-PCM1
GFP-PCM1-N
GFP-PCM1-M
GFP-PCM1
GFP-PCM1-N
GFP-PCM1-M
Total volume of granules (µm³)
Average volume of granules (µm³)
Normalized total granule intensity
Average granule intensity
Granule number
Average distance form centrosome (µm)
Time (min)

**Supplementary Figure S4**

**Spatiotemporal analysis of CS granule properties during mitosis.**

**(A)** Spatiotemporal analysis of *de novo* assembled PCM1 granules during mitosis. Representative still frames from time-lapse confocal images show RPE1::PCM1 KO cells inducibly expressing GFP-PCM1, GFP-PCM1-N, and GFP-PCM1-M. Cells were imaged after the addition of 1 μM doxycycline and SIR-tubulin. Imaging was conducted every 1.5 minute over 144 minutes. Time "0 min" marks the start of granule nucleation. Centrosomes are indicated by cyan arrows at each time point, and cell boundaries are outlined with white dashed lines. Scale bars: 5 μm.

**(B)** Quantification of size, fluorescence intensity and number of PCM1 granules from (D). Average distance to centrosome values were quantified from the distance between granules and SIR-tubulin staining for each cell. Images were taken in 1.5 minute intervals for 144 minutes. Mitosis is shown with faded colors for each quantification. Data represents the mean from quantifications from a cell for each line. Results were consistent with three independent experiments.

### Figure S5

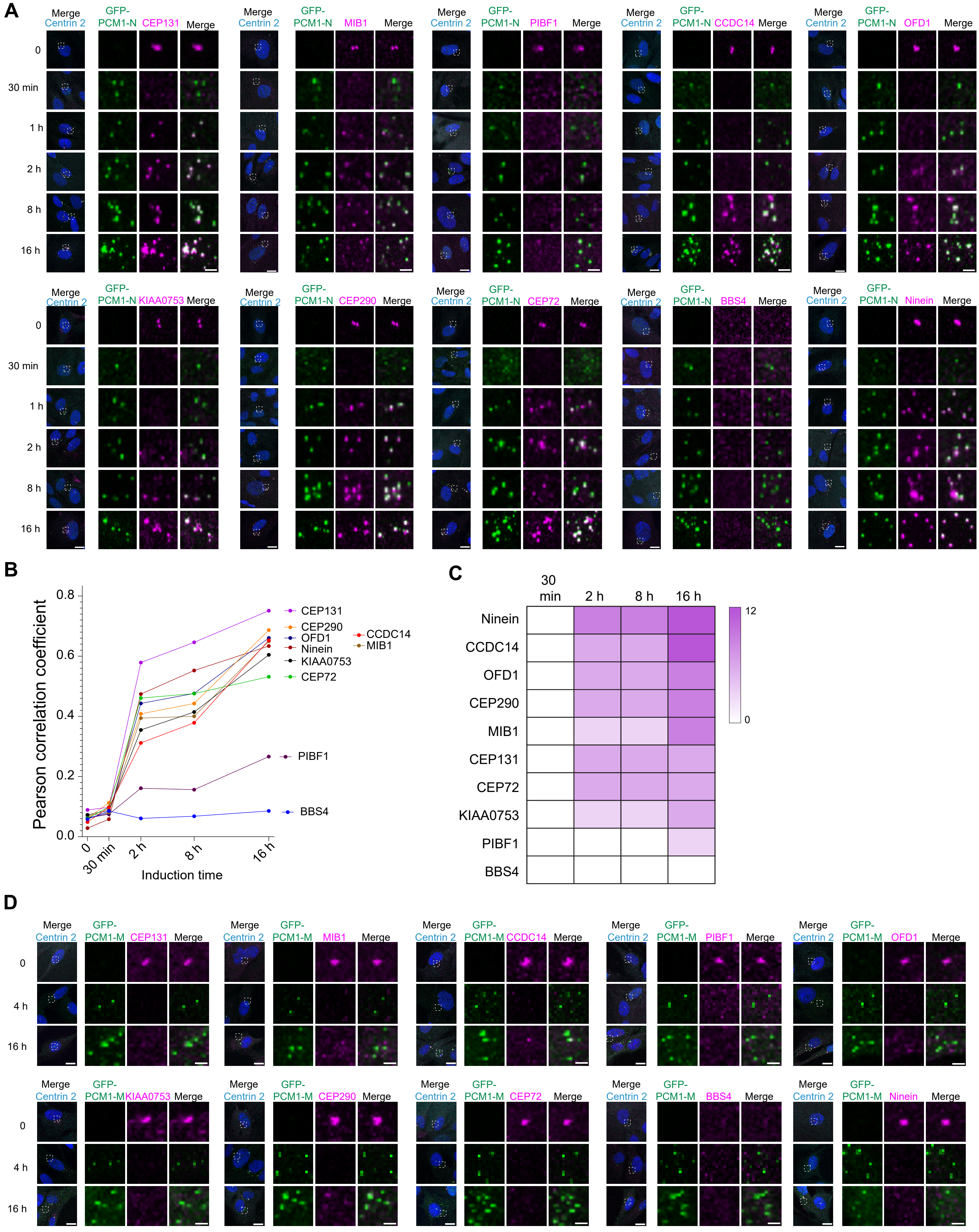
A
Merge
Centrin 2
GFP-PCM1-N
CEP131
MIB1
PIBF1
CCDC14
OFD1
KIAA0753
CEP290
CEP72
BBS4
Ninein
0
30 min
1 h
2 h
8 h
16 h
B
Pearson correlation coefficient
0.0
0.2
0.4
0.6
0.8
0
30 min
2 h
8 h
16 h
Induction time
CEP131
CEP290
OFD1
Ninein
KIAA0753
CEP72
CCDC14
MIB1
PIBF1
BBS4
C
30 min
2 h
8 h
16 h
Ninein
CCDC14
OFD1
CEP290
MIB1
CEP131
CEP72
KIAA0753
PIBF1
BBS4
12
0
D
Merge
Centrin 2
GFP-PCM1-M
CEP131
MIB1
CCDC14
PIBF1
OFD1
KIAA0753
CEP290
CEP72
BBS4
Ninein
0
4 h
16 h

**Supplementary Figure S5**

**Spatiotemporal analysis of client recruitment to scaffolds formed by PCM1-N and PCM1-M during CS biogenesis.**

**(A)** Representative images for co-localization of various CS clients with the PCM1-N scaffold in a *de novo* biogenesis assay. RPE1::PCM1 KO cells inducibly expressing GFP-PCM1-N were fixed before induction and 30 min, 2 hours, 8 hours, 16 hours after induction. They were then stained with antibodies for GFP and clients denoted in magenta. Representative confocal images are shown for each client and time point. White dashed boxes highlight zoomed-in regions. Scale bars: 10 µm (main panels), 2 µm (insets).

**(B)** Pearson correlation coefficient (PCC) analysis of client-scaffold colocalization. PCCs between PCM1-N and client proteins were calculated from 3D images using Imaris Software. They represent colocalization analysis across 30 cells per condition in two independent experiments. The negative control, represented by the PCC between PCM1 and G3BP1 (0.032), indicates the lower detection limit of our colocalization. The positive control, between PCM1 and PCM1 (0.887), represents the upper limit of our colocalization detection.

**(C)** Classification of client recruitment dynamics based on colocalization strength for B. Recruitment time of each client to scaffold PCM1 was determined by comparing Pearson Correlation Coefficient (PCC) values relative to the 0-minute condition. Colocalization strength was classified as positive if the PCC value at 10 minutes was at least twice that at 0 minutes. Darker magenta color represents up-to 5 times increase in the PCC value compared to 0 minutes. These thresholds were established based on the observed colocalization ranges in our dataset. A 2-fold increase marks the minimum detectable enrichment above baseline, ensuring subtle recruitment events are noted, while a 5-fold increase represents the maximum detected, with 3-fold and 3.5-fold serving as intermediate steps. This data-driven approach captures the full spectrum of recruitment behaviors effectively.

**D)** Representative images for co-localization of various CS clients with the PCM1-M scaffold in a *de novo* biogenesis assay. RPE1::PCM1 KO cells inducibly expressing GFP-PCM1-M were fixed before induction and 4 hours and 16 hours after induction. They were then stained with antibodies for GFP and clients denoted in magenta. Representative confocal images are shown for each client and time point. White dashed boxes highlight zoomed-in regions. Scale bars: 10 µm (main panels), 2 µm (insets).

### Figure S6

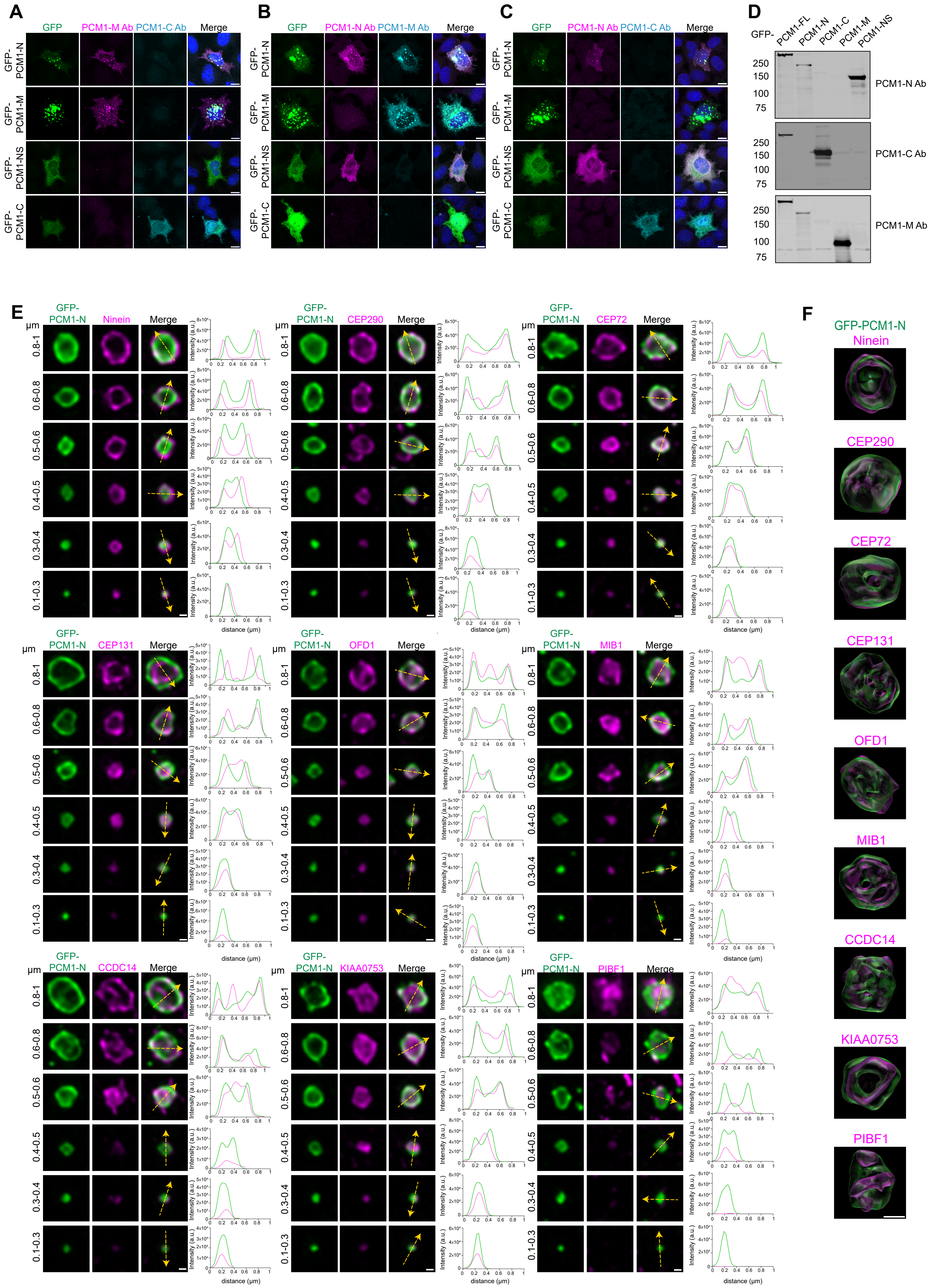
A
GFP
PCM1-M Ab
PCM1-C Ab
Merge
GFP-PCM1-N
GFP-PCM1-M
GFP-PCM1-NS
GFP-PCM1-C
B
PCM1-N Ab
PCM1-M Ab
C
PCM1-N Ab
PCM1-C Ab
D
GFP-
PCM1-FL
PCM1-N
PCM1-C
PCM1-M
PCM1-NS
250
150
100
75
PCM1-N Ab
PCM1-C Ab
PCM1-M Ab
E
GFP-PCM1-N
Ninein
CEP290
CEP72
CEP131
OFD1
MIB1
CCDC14
KIAA0753
PIBF1
μm
0.8-1
0.6-0.8
0.5-0.6
0.4-0.5
0.3-0.4
0.1-0.3
Intensity (a.u.)
distance (μm)
F
GFP-PCM1-N

#### Supplementary Figure S6

**The PCM1 scaffold and its clients are spatially organized into subdomains within CS granules.**

**(A, B, C)** Validation of antibodies targeting distinct regions of PCM1 by immunofluorescence in PCM1 KO cells expressing specific PCM1 fragments. The specificity of PCM1 antibodies for the N-terminal 1-254 aa (PCM1-N-Ab), middle region 700-1200 aa (PCM1-M-Ab), and C-terminus 1664-2024 aa (PCM1-C-Ab) was confirmed through immunofluorescence experiments. Representative confocal images show HeLa::PCM1 KO cells transfected with GFP-PCM1-N, GFP-PCM1-M, GFP-PCM1-NS, and GFP-PCM1-C plasmids for 24 hours. Cells were fixed with 4% PFA and stained for GFP (green) along with various combinations of antibodies: (A) PCM1-M-Ab (magenta), PCM1-C-Ab (cyan); (B) PCM1-N-Ab (magenta), PCM1-M-Ab (cyan); (C) PCM1-N-Ab (magenta), PCM1-C-Ab (cyan). Scale bars: 10 μm

**(D)** Validation of antibodies targeting distinct regions of PCM1 by immunoblotting in PCM1 KO cells expressing specific PCM1 fragments. HEK293T::PCM1 KO cells were transfected with GFP-PCM1-FL, GFP-PCM1-N, GFP-PCM1-C, GFP-PCM1-M, and GFP-PCM1-NS plasmids for 48 hours. Following transfection, cells were lysed, and the samples were subjected to SDS-PAGE and immunoblotting. The samples were then blotted with PCM1-N-Ab, PCM1-C-Ab, or PCM1-M-Ab.

**(E)** Spatial organization of PCM1-N and various CS client proteins revealed by super-resolution microscopy. High-magnification images show granules formed by GFP-PCM1-N colocalizing with different client proteins in RPE1::PCM1-KO cells after doxycycline induction for 24 hours. After fixation, cells were stained with anti-GFP antibody (green) and client proteins (magenta), then imaged using a Zeiss Elyra 7 Lattice SIM super-resolution microscope. Granules with varying diameters, indicated on the left side of each image, were selected as representative. Intensity plot profiles were generated along the yellow arrows for each granule using ImageJ. In the graphs, green represents the fluorescent intensity of PCM1, and magenta represents that of the client proteins. Similar results were obtained from three biological replicates. Scale bars: 0.2 μm.

**(F)** Spatial organization of PCM1-N and various CS clients represented in 3D. Representative images for the spatial organization of PCM1-N and specific client proteins were 3D rendered for enhanced visualization of their relative positions. Scale bar: 0.5 μm.

### Figure S7

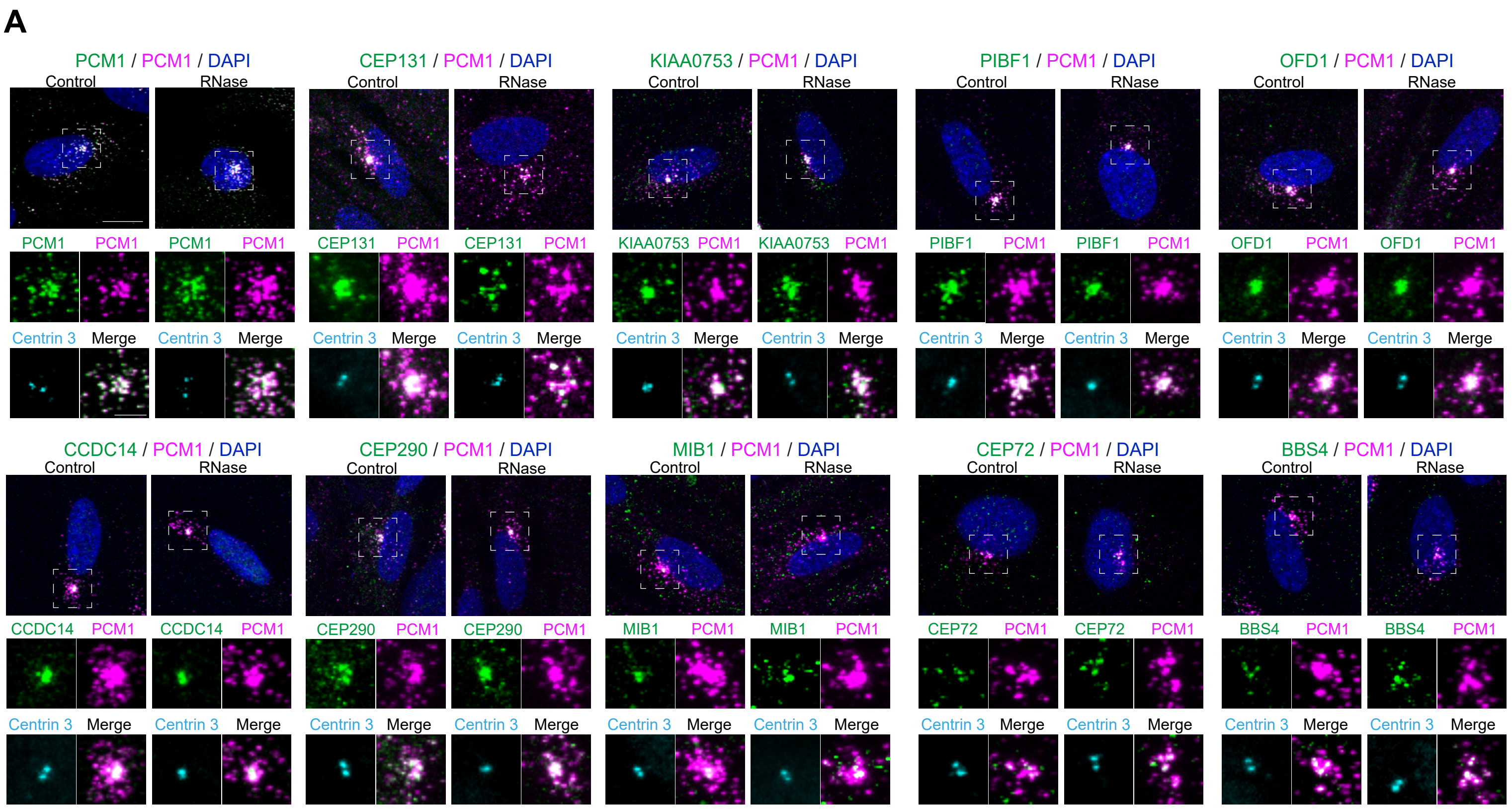

A
PCM1 / PCM1 / DAPI
CEP131 / PCM1 / DAPI
KIAA0753 / PCM1 / DAPI
PIBF1 / PCM1 / DAPI
OFD1 / PCM1 / DAPI
CCDC14 / PCM1 / DAPI
CEP290 / PCM1 / DAPI
MIB1 / PCM1 / DAPI
CEP72 / PCM1 / DAPI
BBS4 / PCM1 / DAPI
Control
RNase
Centrin 3
Merge


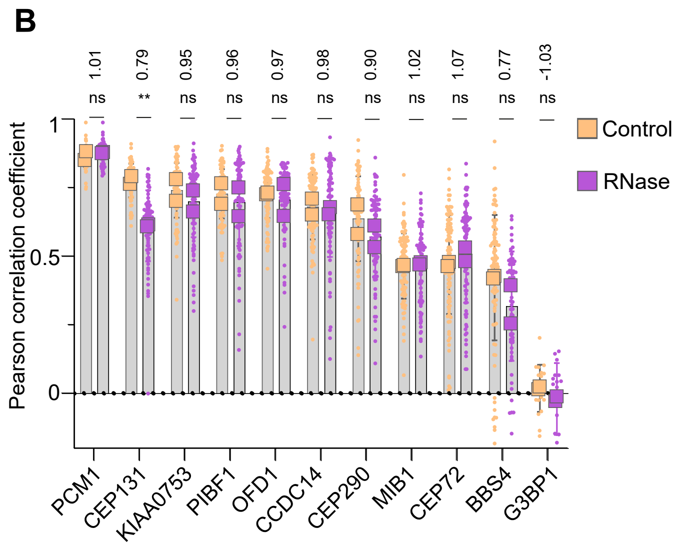

B
1.01
0.79
0.95
0.96
0.97
0.98
0.90
1.02
1.07
0.77
-1.03
ns
**
Pearson correlation coefficient
1
0.5
0
PCM1
CEP131
KIAA0753
PIBF1
OFD1
CCDC14
CEP290
MIB1
CEP72
BBS4
G3BP1
Control
RNase

**Supplementary Figure S7**

**Colocalization index of PCM1 and clients after RNase treatment.**

**(A)** Representative confocal images of co-localization between PCM1 and various clients before and after RNase treatment. RPE1 cells were treated with a combination of 1 mg/ml RNase A and 1 mg/ml RNase I for 1 h, then fixed and stained with antibodies against client proteins (green), PCM1 (magenta), centrin 3 (cyan), and DAPI (blue). White boxes indicate zoomed-in regions. Scale bars: 10 μm (main panels), 5 μm (insets).

**(B)** Pearson correlation coefficient quantifying colocalization between PCM1 and client proteins. Fold change is calculated by comparing the mean value of RNase-treated samples to that of the control. Data are from 40 cells per replicate, with two independent experiments. Statistical significance was determined using a t-test comparing treatment and control for each colocalization pair (ns > 0.05, **P < 0.01).

### Figure S8

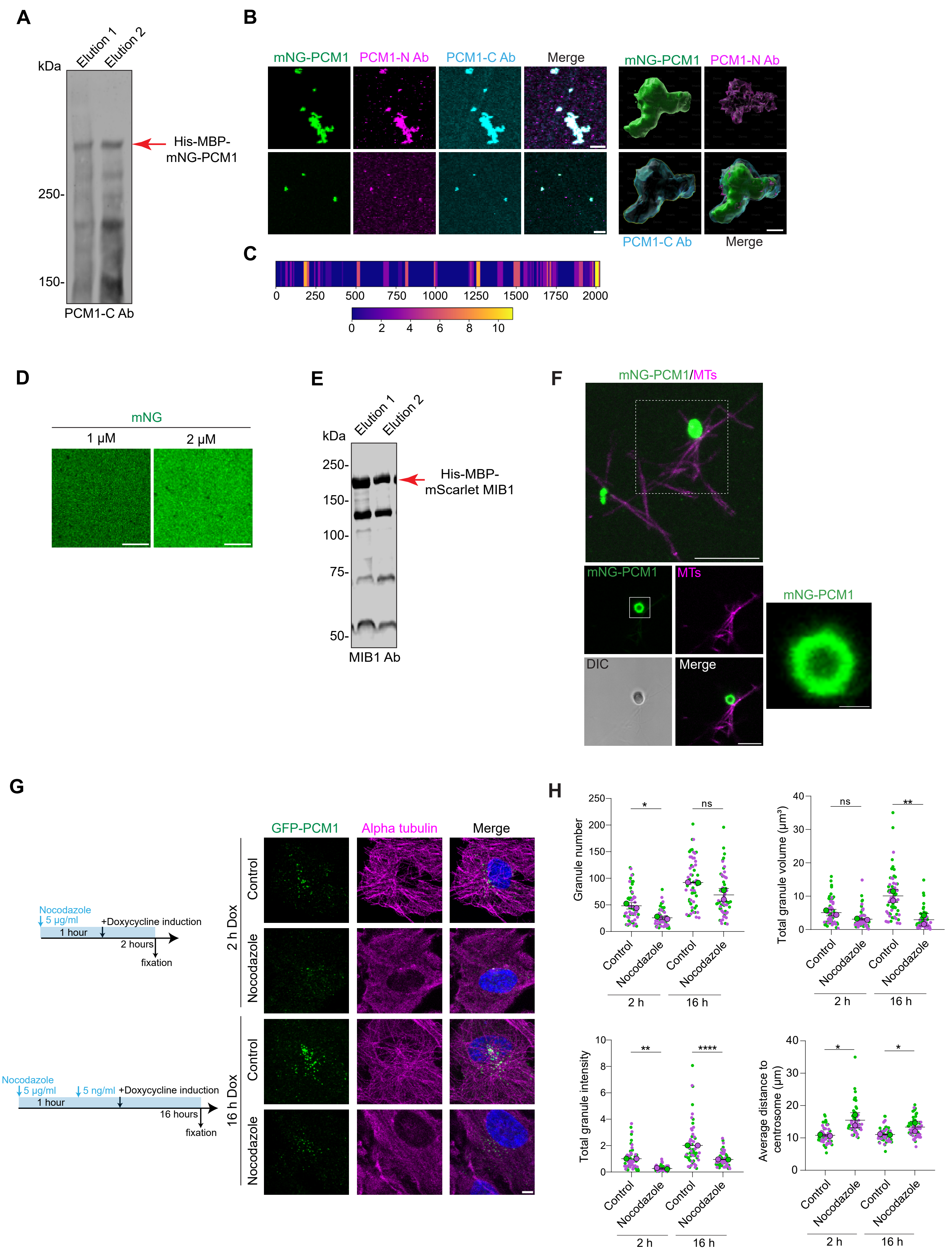

A
kDa
Elution 1
Elution 2
His-MBP-mNG-PCM1
250-
150-
PCM1-C Ab
B
mNG-PCM1
PCM1-N Ab
PCM1-C Ab
Merge
C
0
250
500
750
1000
1250
1500
1750
2000
0
2
4
6
8
10
D
mNG
1 μM
2 μM
E
kDa
250-
150-
100-
75-
50-
His-MBP-mScarlet MIB1
MIB1 Ab
F
mNG-PCM1/MTs
MTs
DIC
Merge
G
Nocodazole
5 μg/ml
+Doxycycline induction
1 hour
2 hours
fixation
5 ng/ml
16 hours
GFP-PCM1
Alpha tubulin
Control
Nocodazole
2 h Dox
16 h Dox
H
Granule number
Total granule volume (μm³)
Total granule intensity
Average distance to centrosome (μm)
ns
2 h
16 h

**Supplementary Figure S8**

**Characterization of biophysical properties of CS.**

**(A)** Confirmation of His-MBP-mNG-PCM1 purification. Analysis of purified mNeonGreen-PCM1 separated by SDS-PAGE and immunoblotted with PCM1-C Ab. Arrow indicates full-length PCM1 band.

**(B)** Confirmation of His-MBP-mNG-PCM1 purification by immunofluorescence. mNeonGreen-PCM1 condensates in phase separation buffer were fixed with 15 % glutaraldehyde, spun on cover slips, fixed with 4% PFA and stained for PCM1 N-term and C-term specific antibodies. 3D images were exported from Imaris software. Scale bar: 5 μm.

**(C)** Confirmation of His-MBP-mNG-PCM1 purification by mass spectrometry. Mass spectrometry analysis of purified mNeonGreen-PCM1 for peptide coverage marked on the 2024 aa long human protein sequence. The negative gradient of purple to yellow indicates peptide regions that have been identified 0-10 times on the sequence where purple indicates 0 and yellow indicates 10.

**(D)** Analysis of His-MBP-mNG tag as a control in phase separation experiments. Purified His-MBP-mNG in phase separation buffer at concentrations 1 and 2 μM observed as soluble protein for control of specificity of PCM1-mediated phase separation. Scale bar: 10 μm.

**(E)** Confirmation of His-MBP-mScarlet MIB1 purification. Analysis of purified mScarlet-MIB1 separated by SDS-PAGE and immunoblotted with MIB1 Ab. Arrow indicates full-length MIB1 band.

**(F)** mNeonGreen-PCM1, and taxol stabilized microtubules polymerized from purified bovine tubulin labeled with Alexa Fluor 568 were set in BRB80 buffer, flown in chambers and imaged. Scale bar, 10 μM for field of view at single z-stack; 2 μM for inset. mNeonGreen-PCM1 and taxol-stabilized microtubules, polymerized from purified bovine tubulin labeled with Alexa Fluor 568, were prepared in BRB80 buffer, introduced into chambers, and imaged. Insets represent specific z-stacks to highlight the spatial organization of PCM1-N. Scale bars: 10 μm (main panels), 2 μm (insets).

**(G)** Microtubules promote CS assembly. Representative confocal images of RPE1::PCM1 KO cells inducibly expressing GFP-PCM1 treated with nocodazole. Cells were either treated with 5 μg/ml nocodazole for 2 hours before 1 μM doxycycline induction for 2 hours or with 5 μg/ml nocodazole for 1 hour, then with 5 ng/ml nocodazole and 1 μM doxycycline for 16 hours. Results are from two biological replicates, n=30 cells per replicate. Cells were fixed and stained for anti-GFP (green), anti-alpha tubulin 2 (magenta) and DAPI for nucleus (blue). Scale bar: 2 μm.

**(H)** Quantification of CS granule properties after nocodazole treatment from (D). Granule properties, including total volume of granules per cell ($\mu m^3$), average volume of granules per cell ($\mu m^3$), total granule intensity, granule number and average distance of granules to centrosome (μm) values were measured using Imaris software. Centrosomes were identified using centrin 2 staining to quantify the distance values. Statistical significance was determined by the Student's t-test (paired and two-tailed). Results are from two biological replicates, green and magenta colored dots representing different replicates. n=30 cells cells per replicate. *:P<0.05, **: P<0.01, ***:P<0.001, ****:P<0.0001; ns: not significant.
